# Dissimilar conservation pattern in hepatitis C virus mutant spectra, consensus sequences, and data banks

**DOI:** 10.1101/2020.07.03.186171

**Authors:** Carlos García-Crespo, María Eugenia Soria, Isabel Gallego, Ana Isabel de Ávila, Brenda Martínez-González, Lucía Vázquez-Sirvent, Jordi Gómez, Carlos Briones, Josep Gregori, Josep Quer, Celia Perales, Esteban Domingo

**Affiliations:** Department of Interactions with the environment, Centro de Biología Molecular “Severo Ochoa” (CSIC-UAM), Consejo Superior de Investigaciones Científicas (CSIC), Campus de Cantoblanco, 28049, Madrid, Spain; Department of Clinical Microbiology, IIS-Fundación Jiménez Díaz, UAM. Av. Reyes Católicos 2, 28040 Madrid, Spain; Centro de Investigación Biomédica en Red de Enfermedades Hepáticas y Digestivas (CIBERehd) del Instituto de Salud Carlos III, 28029, Madrid, Spain; Department of Molecular Biology, Instituto de Parasitología y Biomedicina López-Neyra’ (CSIC), Parque Tecnológico Ciencias de la Salud, Armilla, 18016, Granada, Spain; Department of Molecular Evolution, Centro de Astrobiología (CAB, CSIC-INTA), 28850 Torrejón de Ardoz, Madrid, Spain; Liver Unit, Internal Medicine Hospital Universitari Vall d’Hebron, Vall d’Hebron Institut de Recerca (VHIR), 08035, Barcelona, Spain; Roche Diagnostics, S.L., Sant Cugat del Vallés, 08174, Barcelona, Spain

**Keywords:** antiviral intervention, consensus sequence, mutant spectrum, residue conservation, viral ligands, virus data banks

## Abstract

The influence of quasispecies dynamics on long-term virus diversification in nature is a largely unexplored question. Specifically, whether intra-host nucleotide and amino acid variation in quasispecies fits variation observed in consensus sequences or data bank alignments is unknown. Genome conservation and dynamics simulations are used for the computational design of universal vaccines, therapeutic antibodies and pan-genomic antiviral agents. The expectation is that selection of escape mutants will be limited when mutations at conserved residues are required. This strategy assumes long-term (epidemiologically relevant) conservation but, critically, does not consider short-term (quasispecies-dictated) residue conservation. We have calculated mutant frequencies of individual loci from mutant spectra of hepatitis C virus (HCV) populations passaged in cell culture and from infected patients. Nucleotide or amino acid conservation in consensus sequences of the same populations, or in the Los Alamos HCV data bank did not match residue conservation in mutant spectra. The results relativize the concept of sequence conservation in viral genetics, and suggest that residue invariance in data banks is an insufficient basis for the design of universal viral ligands for clinical purposes. Our calculations suggest relaxed mutational restrictions during quasispecies dynamics, which may contribute to higher calculated short-term than long-term viral evolutionary rates.

## Introduction

Hepatitis C virus (HCV) is an important human pathogen that poses many global public health challenges, including the lack of a vaccine, and a non-universal accessibility of effective treatments [1]. We are interested in its quasispecies dynamics since it has proven pertinent to the understanding of HCV-associated liver disease progression, and treatment efficacy [2–4] and its possible influence on virus evolution at the epidemiological level is an open question [5–7]. In particular, it is not known if long-term residue conservation as calculated from viral genome alignments of consensus sequences in data banks corresponds to limited variation of the same residues in mutant spectra. A current aim in the control of diseases associated with RNA viruses is the computer- and dynamics simulations-guided design of universal vaccines, therapeutic antibodies or pan-genomic antiviral agents based on conserved viral residues, with conservation defined according to the alignment of nucleotide and amino acid sequences recorded in data banks. The expectation is to render reagents effective against all (or a majority of) circulating serotypes and genotypes of a virus [8–13]. This strategy rests on the assumption that viral genomic residues which are conserved among independent viral isolates tend to be also conserved within the mutant spectra displayed by the viruses during their intra-host multiplication. Yet, it will be during such stage of the infection cycle when the binding ligands should prevent viral progeny production, or when the newly assembled and released viral particles and newly infected cells should be controlled by a vaccine-induced immune response. The critical assumption of parallel residue conservation in mutant spectra and data banks has not been tested. This point is relevant because it interrogates a relationship between two ranks of operation of virus evolution: one which is associated with intra-host, short-term mutant cloud dynamics, and another that involves long-term evolution at the epidemiological level, generally evaluated with consensus sequences.

Here we describe a two-step approach to this question with HCV. First, we compare the degree of individual residue conservation (percentage of residue identity) in the course of viral quasispecies dynamics —quantified by the mutant frequency of each variable residue in cell culture and infected patients— with the conservation deduced from an alignments of the consensus sequences of the same populations. Second, we compare conservation levels in mutant spectra with those of the same residues in the Los Alamos National laboratory (LANL) data bank sequence alignment. Mutant frequencies in quasispecies have been obtained in cell culture thanks to the availability of infectious molecular clones of HCV that permit experimental evolution designs with this important pathogen [14–18]. In a recent study we examined mutant spectrum variations upon subjecting a clonal HCV population to 200 serial infections of Huh-7.5 cells, using fresh cells at each passage, thereby preventing co-evolution of the host cells [19]. Under these conditions, a variation of mutant frequency of several individual genomic residues even between successive passages was consistently observed in three biological replicas. Because mutant frequency variations were continuous, we termed them “mutational waves” to emphasize a difference with those residues whose frequency remained constant with passage number [17,19]. Mutational waves were characterized by deep sequencing analysis of 1,005 genomic positions from the NS5A-NS5B-coding region, and they involved 145 different mutations. The dependability of the deep sequencing analyses was sustained on experimental and bioinformatics controls that yielded reliable cut-off mutant frequency values of 0.5% [19–21] (see also Materials and Methods). Furthermore, Sanger sequencing of the HCV p0, HCV p100, HCV p200 and their derivatives at passage 4 identified 114 heterogeneous sites (two different nucleotides at the same genomic position) within the NS2- to NS5B-coding region, thus confirming mutant frequency variations by two independent procedures [19].

In addition, we have included in our study 522 different amino acid substitutions in the NS5B mutant spectra of a cohort of 220 HCV-infected patients that failed antiviral therapy [22]. This information on HCV quasispecies composition in cell culture and *in vivo* has documented that residue variation at the quasispecies level does not match conservation in the corresponding consensus sequences or in the sequences deposited in the LANL data bank. The computations suggest a remarkably higher mutant tolerance in evolving HCV quasispecies than reflected in consensus sequences. This difference may contribute also to a higher calculated intra-host than inter-host evolutionary rates, an intriguing difference observed with several viral pathogens [6,7,23–27]. A new residue conservation score can be produced upon analysis of sequence alignments of the genomes in mutant spectra that may be more appropriate for the design of universal ligands or vaccines intended to control RNA viruses.

## Results

### HCV residues involved in mutational waves and their conservation in the consensus sequences of the populations

Two-hundred serial passages of a clonal [derived by transcription of plasmid Jc1FLAG2(p7-nsGLuc2A) (abbreviated Jc1Luc)] HCV population [28] in Huh-7.5 cells identified 145 different mutations —located within genomic nucleotides 7649 to 8653— that participated in mutational waves based on deep sequencing data of 39 HCV populations [19] (Fig. 1A) [residue numbers are according to HCV reference isolate JFH-1 (accession number #AB047639); mutations and deduced amino acid substitutions are given in Table S1 (http://babia.cbm.uam.es/~lab121/SupplMatGarcia-Crespo)]. The alignment of the consensus nucleotide sequences for each of the 39 populations indicated 98% sequence identity. The residues that participated in mutational waves were scattered along the region analyzed, with an accumulation within residues 7650 to 7661 (those encoding the C-terminal amino acids of NS5A and the N-terminal amino acids of NS5B) (Fig. 2). A similar distribution was obtained at the amino acid level (Fig. S1). To assign each variable residue in the quasispecies to a degree of conservation according to the consensus sequence alignment, the 1005 nucleotides comprised between positions 7649 and 8653 were divided into ten conservation categories calculated relative to the most abundant nucleotide at the corresponding position in the alignment. A residue that falls in the 90%-100% conservation window means that the residue is present in at least 90% of the consensus sequences; this is followed by lower conservation categories, with 20%-30% as the minimum possible conservation window at the nucleotide level. The 145 nucleotides (and deduced amino acids) that varied in frequency in the HCV quasispecies belonged to high conservation categories in the consensus sequence alignment (Fig. 3A,B). The difference between the observed residue distributions and those in which an equal number of mutations was assigned uniformly among the possible conservation groups was statistically significant both for nucleotides and amino acids (p=0.0004998 and p=0.0004998, respectively; Pearson’s chi-squared test with Monte Carlo correction). To rectify a possible bias derived from the great abundance of residues in the 90%-100% conservation window, values were normalization to the number of positions that fall in its conservation group. This correction shifted the maximum number of residues from the 90%-100% to the 80%-90% window (Fig. 3C,D). In this case, the difference between the observed distribution and one in which the number of mutations was equally distributed among the possible conservation groups was not statistically significant (p= 1 for nucleotides and amino acids; Pearson’s chi-squared test with Monte Carlo correction). Conversion of complex mutant spectra into consensus sequences determines a ranking of residue conservation that does not correspond with conservation at the level of the mutant spectra that yielded the consensus sequences.

**Figure 1.**
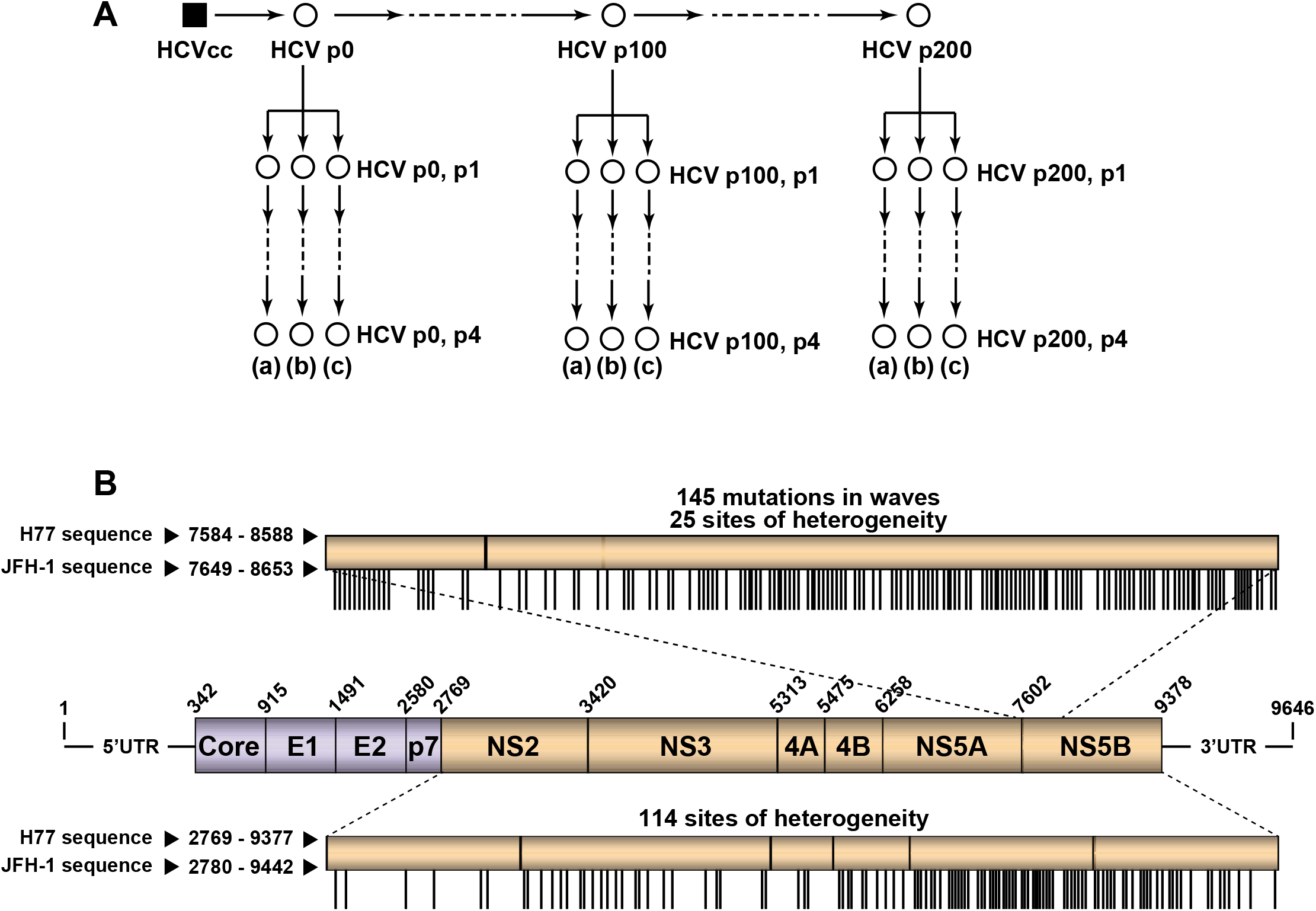
Experimental design, and HCV genome analysis. **(A)** Clonal HCV population HCVcc prepared by transcription of plasmid Jc1Luc [28], followed by RNA electroporation into Huh-7 Lunet cells, was amplified to produce HCV p0, and the population subjected to 200 passages in Huh-7.5 cells, as described [17–19]. The initial population HCV p0 and the populations at passages 100 (HCV p100) and 200 (HCV p200) were passaged four additional times in Huh-7.5 cells in triplicate [replicas (a), (b), (c)]. The mutant spectrum of each of the populations (represented by empty circles) was analyzed. **(B)** Scheme of the HCV genome, encoded proteins and genomic regions analyzed; residue numbering is according to isolate H77. In the boxes depicted above and below the genome, the equivalence of nucleotide residue numbers for isolates H77 and JFH-1 is indicated. The upper box corresponding the NS5A-NS5B-coding region indicates the positions involved in mutational waves within residues 7584 to 8588 (H77 numbering) (horizontal lines). Frequency levels are described in Materials and Methods and indicated in Table S1 (http://babia.cbm.uam.es/~lab121/SupplMatGarcia-Crespo). The bottom box corresponding to the NS2-NS5B-coding region indicates the sites of heterogeneity determined by Sanger sequencing (see also Materials and Methods). Results are from [19], and the mutations under study are compiled in Tables S1 and S3 (http://babia.cbm.uam.es/~lab121/SupplMatGarcia-Crespo).

**Figure 2.**
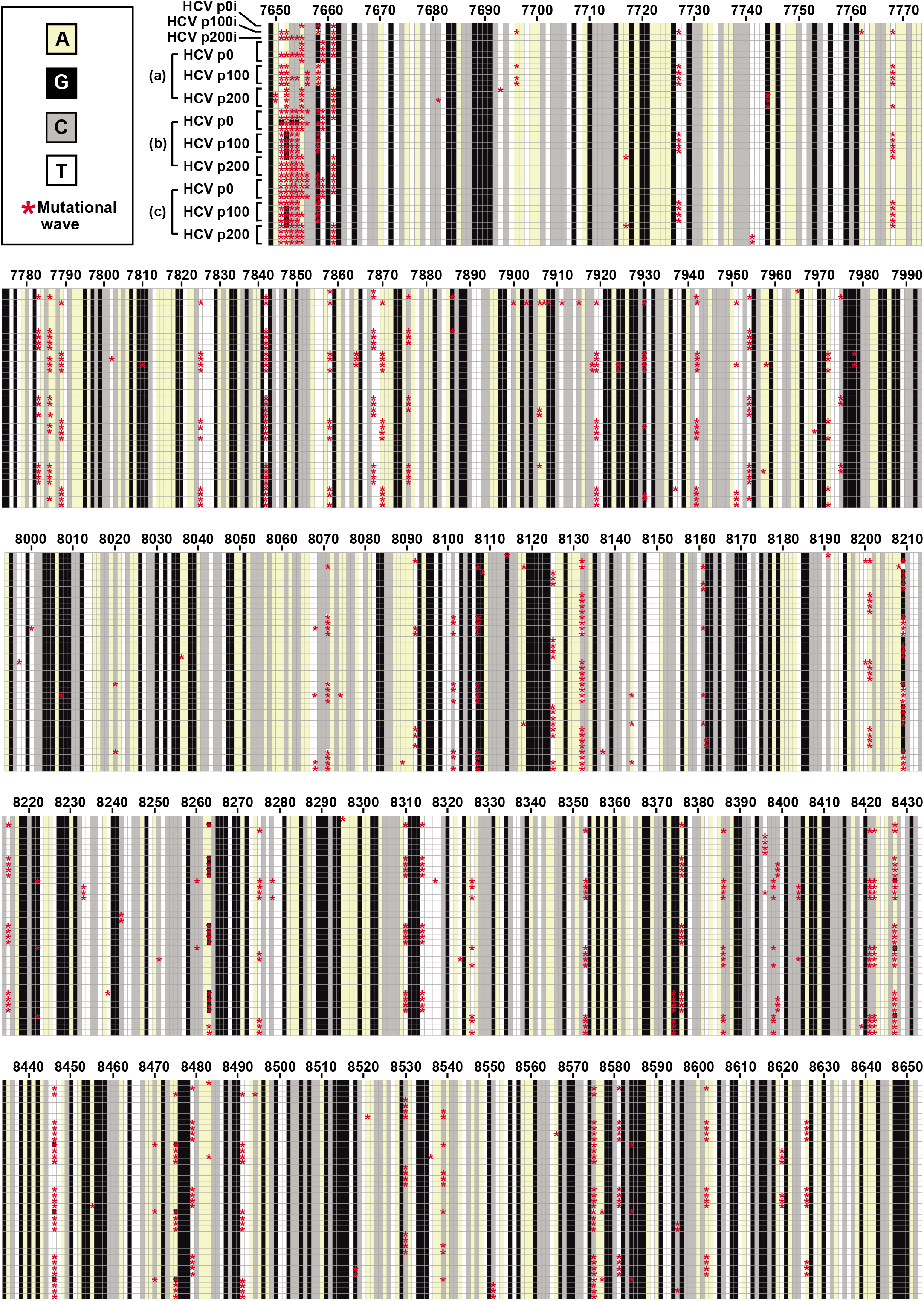
Heat map of consensus sequences of 39 HCV populations derived from HCV p0 upon passage in Huh-7.5 cells. The region analyzed by deep sequencing spans genomic residues 7649 to 8653 (residue numbering according to reference isolate JFH-1). The populations are those identified by empty circles in the HCV passage diagram of Fig. 1A, and are indicated at the left of the top bloc of the alignment; (a), (b), and (c) refer to the three replicas of the four serial passages of HCV p0, HCV p100 and HCV p200. Each horizontal alignment of squares displays the consensus sequence (1005 nucleotide positions) of the population written on the left, with the nucleotide color code given in the upper box on the left. The red asterisks indicate the nucleotides that participated in mutational waves (that is, that changed in frequency among any of the populations analyzed). The complete set of mutations and deduced amino acid substitutions are given in Table S1 (http://babia.cbm.uam.es/~lab121/SupplMatGarcia-Crespo).

**Figure 3.**
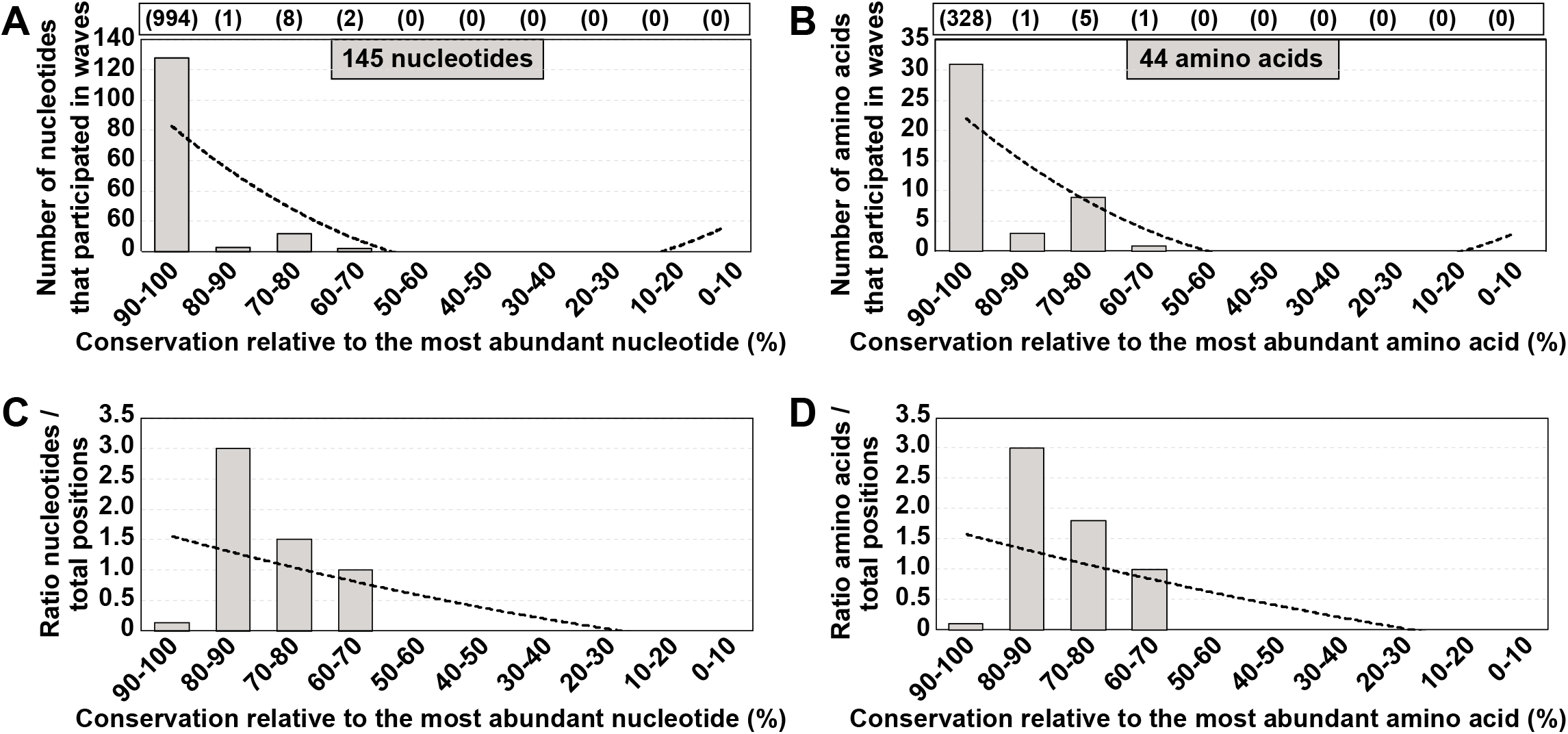
Degree of conservation of those residues that participated in mutational waves. **(A)** Number of nucleotides involved in mutational waves distributed among conservation groups, calculated relative to the most abundant nucleotide at the corresponding position in the alignment of 39 consensus sequences. Conservation groups are indicated in abscissa, and the number of nucleotides that participated in mutational waves in each group is given in ordinate. The total number of nucleotides within residues 7649 to 8653 from the alignment that fall in each conservation category is indicated in parenthesis in the upper box. The discontinuous line corresponds to function y=2.8636x^2^-39.009x+118.8 (R^2^=0.6221). **(B)** Same as A but at the amino acid level. The discontinuous line corresponds to function y = 0.6818x^2^ – 9.6091x + 31 (R^2^ = 0.7135). **(C)** Data of A normalized to the number of residues in each conservation group; normalization was done by dividing the latter number by the total number of residues from the consensus sequence alignment that fell into the corresponding group. The discontinuous line corresponds to function y = 0.0086x^2^ -0.2926x + 1.8409 (R2 = 0.3595). **(D)** Data of B normalized to the number of residues in each conservation group. The discontinuous line corresponds to function y = 0.0067x^2^-0.2788x+1.8651 (R2 = 0.3577). The position of each mutation and amino acid substitution is given in Table S1 (http://babia.cbm.uam.es/~lab121/SupplMatGarcia-Crespo).

### Conservation in mutant spectra as compared with conservation in the Los Alamos data bank

Given that sites of variability in mutant spectra of the experimental populations did not match variability in consensus sequences, we explored whether the same discrepancy was observed when mutant spectra were confronted with residue conservation in the HCV sequence repository in the LANL data bank. The interest in this comparison resides in the fact that LANL alignments are often taken as a reference for residue conservation in viral genomes, and taken into consideration for the design of broad spectrum viral ligands or vaccines [8–13]. To this aim, a total of 1191 HCV genomic sequences of LANL (https://hcv.lanl.gov/content/sequence/HCV/ToolsOutline.html) were used to define conservation groups, following the same procedure as for the consensus sequences of the HCV quasispecies in cell culture. Inclusion criteria and subtype distribution are given in Materials and Methods, and the accession numbers are listed in Table S2 (http://babia.cbm.uam.es/~lab121/SupplMatGarcia-Crespo); 95.4% of the sequences are from infected patients (of which 26.1% are from antiviral-treated patients, 55.9% from untreated patients, and 17.8% without treatment information). The degree of conservation was calculated for residues 7584 to 8588 of sequence H77 [GenBank accession number AF009606, which is used as reference in the LANL alignment [29]; residues 7584 to 8588 of H77 are equivalent to residues 7649 to 8653 of JFH-1]. Conservation groups were calculated relative to the most abundant nucleotide and amino acid at the corresponding position in the LANL alignment. Then, the position of each of the 145 nucleotides (and of 44 deduced amino acids) that were involved in mutational waves was assigned to the conservation group of the same position in the alignment. The wave mutations were distributed between the 90%-100% and 30%-40% conservation range, with 54.5% of wave mutations falling into the 80%-100% conservation groups (Fig. 4A). The difference between this distribution and one in which an equal number of mutations was assigned uniformly among the possible conservation groups was statistically significant (p=2.8113x10^-9^, chi-square test).

**Figure 4.**
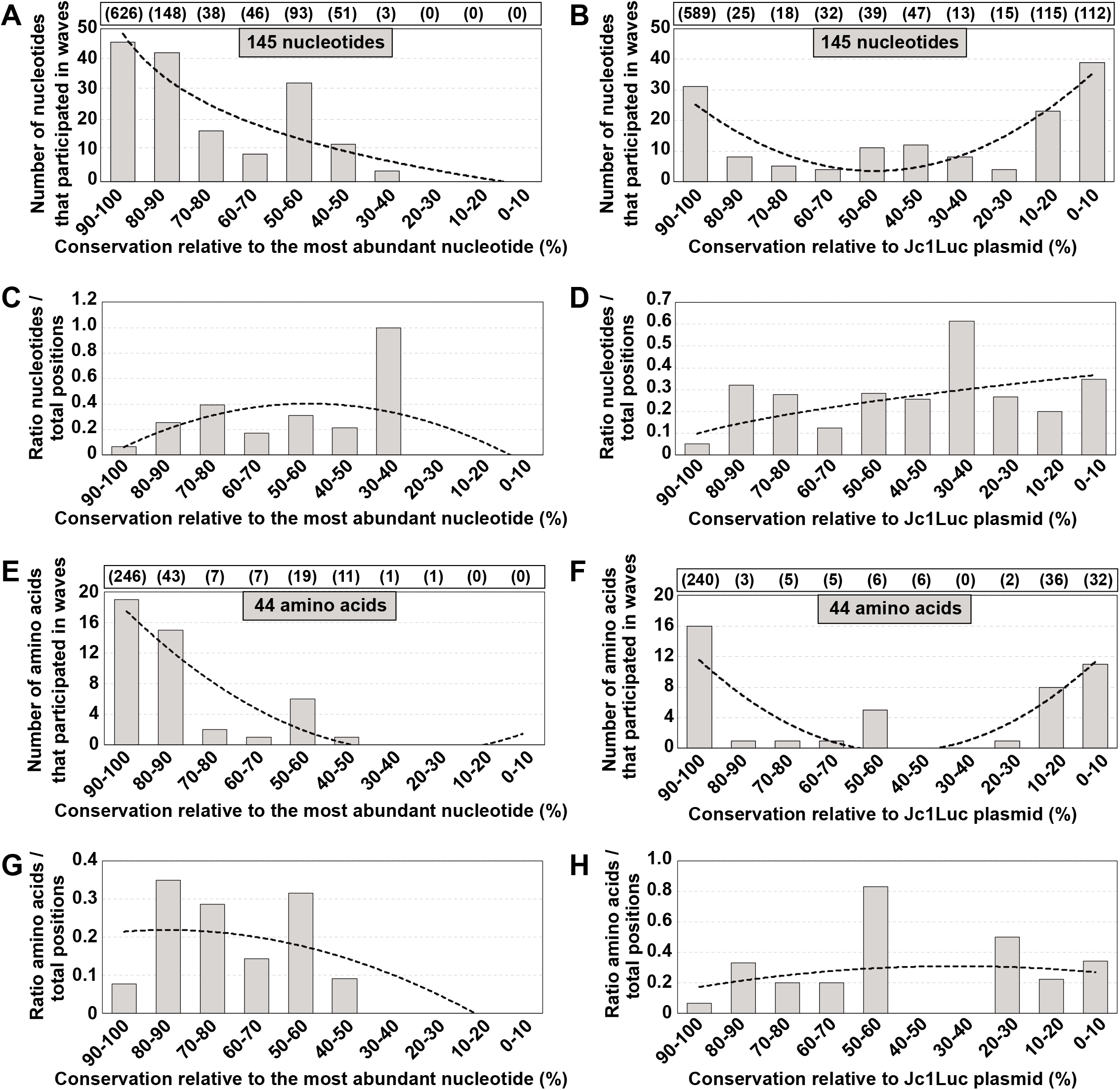
Degree of conservation of nucleotides and amino acids that participated in mutational waves, according to the LANL alignment. **(A)** Number of nucleotides involved in mutational waves distributed among conservation groups, calculated relative to the most abundant nucleotide at the corresponding position in the LANL alignment. Conservation groups are indicated in abscissa, and the number of nucleotides that participated in mutational waves in each group is given in ordinate. The total number of nucleotides within residues 7584 to 8588 (H77 numbering) from the alignment that fall in each conservation category is indicated in parenthesis in the upper box. The discontinuous line corresponds to function y = -19.21ln(x) + 43.511 (R2 = 0.7816). **(B)** Same as A but with nucleotide conservation in the LALN alignment calculated relative to the corresponding residues in plasmid Jc1Luc [28]. The discontinuous line corresponds to function y = 1.3068x^2^ – 13.254x + 37.083 (R2 = 0.7401). **(C)** Data of A normalized to the number of residues in each conservation group; normalization was done by dividing the latter number by the total number of residues from the LANL alignment that fell into the corresponding group. The discontinuous line corresponds to function y = -0.0194x^2^ + 0.2013x – 0.1189 (R2 = 0.2582). **(D)** Data of B normalized to the number of residues in each conservation group. The discontinuous line corresponds to function y = 0.0989x0.5688 (R2 = 0.3987). **(E-H)** Same as A-D but at the amino acid level. The defining functions are E: y = 0.4205x^2^ – 6.4068x + 23.45 (R2 = 0.8178); F: y = 0.5871x^2^ – 6.4826x + 17.45 (R2 = 0.6587); G: y = -0.0045x^2^ + 0.0175x + 0.2018 (R2 = 0.5213); H: y = 0.0644ln(x) + 0.1726 (R^2^ = 0.0347). The position of each mutation and amino acid substitution is given in Table S1, and control calculations and simulations to assess the statistical relevance of the mutant distributions are described in Fig. S3 and Table S6 (http://babia.cbm.uam.es/~lab121/SupplMatGarcia-Crespo).

Given the absence of mutational wave residues in the low conservation categories, we wanted to exclude the possibility that the reference chosen for the calculation may have biased the mutation sites towards high conservation groups. Therefore, we also calculated the conservation ranges in the LANL alignment using as reference the corresponding residues in the HCV sequence in plasmid Jc1Luc. Since the sequence belongs to the specific HCV subtype 2a, and the LANL alignment includes HCV genomes of different subtypes, the reference modification was expected to increase the number of positions belonging to the low conservation windows. With this reference, the plot of the number of nucleotides or amino acids in each conservation category yielded a U-shaped curve (Fig. 4B). A considerable proportion (46%) of wave-involved residues belonged to the two extreme 90-100% or 0-10% conservation groups. Normalization to the total number of sites in each conservation category resulted in a broader distribution across the conservation spectrum (Fig. 4C,D), with no significant difference with a uniform distribution among conservation groups (p>0.999, chi-square test with Monte Carlo correction). Similar results were obtained at the amino acid level (Fig. 4E-H). Thus, there is no correlation between sites of variation in mutational waves of HCV evolving in cell culture and sites of low residue conservation in the LANL alignment.

### Extension of the calculations to mutations associated with sites of heterogeneity in consensus sequences

An independent evaluation of individual mutant spectrum residues that vary in frequency in HCV evolving in cell culture was provided by the genomic positions that contain more than one nucleotide, as quantified in consensus sequences determined by Sanger sequencing for HCV p0, HCV p100, HCV p200, and these populations at passage 4. A total of 25 points of heterogeneity were scored within residues 7584 to 8588 (H77 numbering) of populations HCV p0, HCV p100 and HCV p200 [19] (Fig. 1B and Table S3 in http://babia.cbm.uam.es/~lab121/SupplMatGarcia-Crespo). A plot of the nucleotides at sites displaying heterogeneity versus the degree of conservation of those nucleotide and amino acid sites in the consensus sequences of the cell culture HCV populations or in the LANL alignment, confirmed the results obtained with the wave mutations (Fig. S2) in http://babia.cbm.uam.es/~lab121/SupplMatGarcia-Crespo). Again, residues which vary in mutant frequency —as determined by a procedure which is independent of deep sequencing— during quasispecies dynamics were not the most variable in consensus sequences from progeny HCV p0 populations or those reported in the LANL alignment.

### Extension to other HCV genomic regions

To exclude that the NS5A-NS5B residues analyzed might display a behavior which is not representative of other HCV genomic regions, we extended the calculations to a total of the 114 genomic sites that display composition heterogeneity, located between nucleotides 2769 (beginning of the NS2-coding region) and 9377 (end of the NS5B-coding region) (H77 numbering) of populations HCV p0, HCV p100 and HCV p200 [19] (Fig. 1B and Table S3 in http://babia.cbm.uam.es/~lab121/SupplMatGarcia-Crespo). The results (Fig. 5) are very similar to those obtained with the NS5A-NS5B region. The difference between the distribution of the 114 nucleotides and one in which the distribution included an equal number of sites across the different possible conservation groups, was statistically significant (p=1.0371x10^-7^; chi-square test). In conclusion, residue invariance (undetectable mutant frequencies by current deep sequencing and Sanger sequencing methodology) in mutant spectra of HCV evolving in Huh-7.5 cells does not correlate with conservation of the same residues in consensus sequences or among epidemiologically distant isolates as represented in LANL.

**Figure 5.**
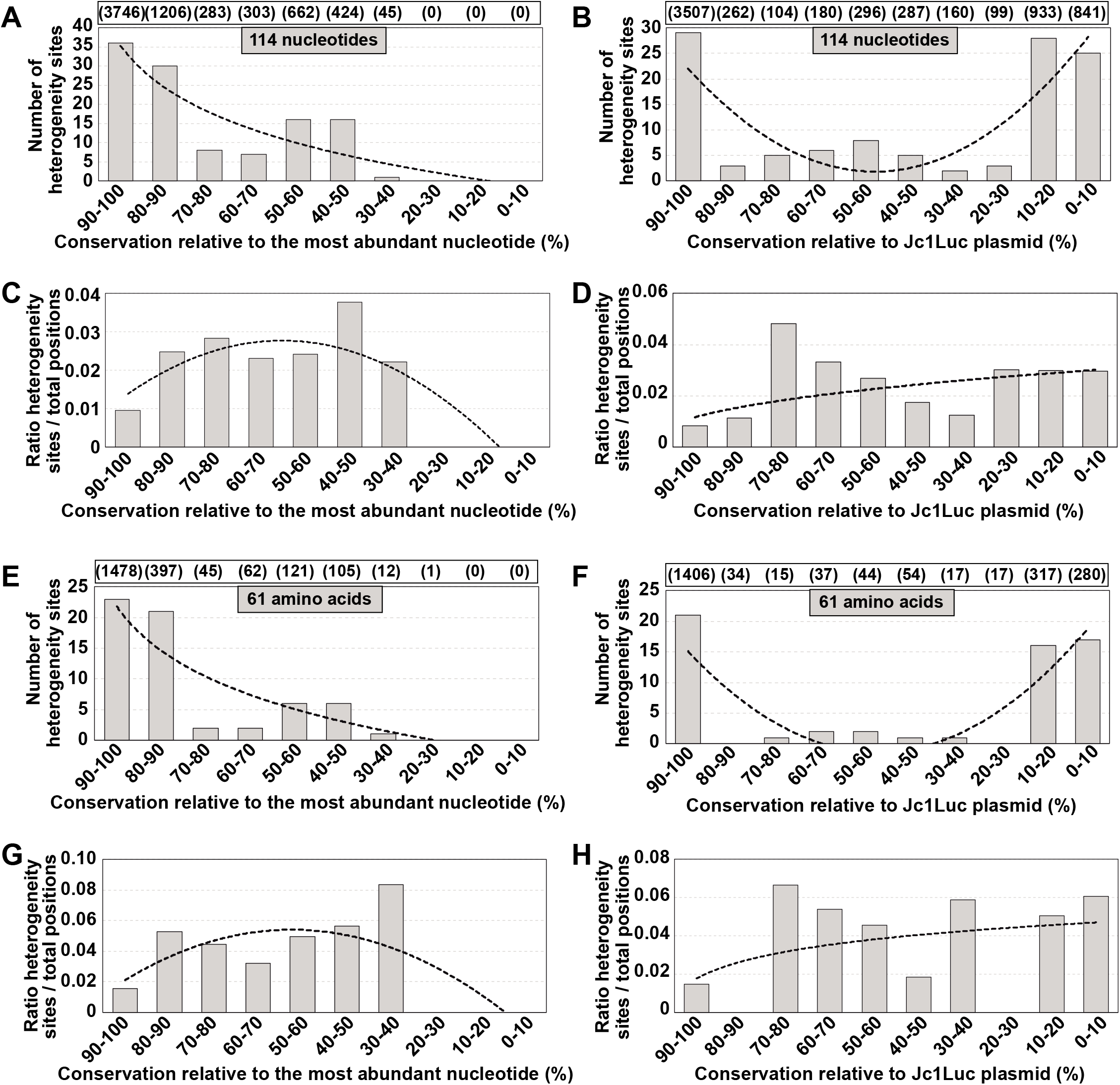
Distribution of sites of heterogeneity in HCV quasispecies among residue conservation groups defined according to LANL. Sites of heterogeneity are those identified within HCV genomic residues 2769 and 9377 (H77 numbering). In this case the analysis was performed separately with 983 sequences of genotype G1 and 129 sequences of genotype G2 [excluding one G2c and one G2k sequence due to the presence of an insertion (https://hcv.lanl.gov/content/sequence/HCV/ToolsOutline.html)]. **(A)** The genomic position at which each heterogeneity site is found is assigned to the conservation range calculated relative to the most abundant nucleotide in each position according to the LANL alignment. Conservation groups are indicated in abscissa, and the number of heterogeneity sites in each group is given in ordinate. The total number of nucleotides that fall in each conservation category is indicated in parenthesis in the upper box. The discontinuous line corresponds to function y = -15.83ln(x) + 35.306 (R2 = 0.7957). **(B)** Same as A except that the conservation groups in LANL alignment were calculated using the HCV sequence in plasmid Jc1Luc as reference. The discontinuous line corresponds to function y = 1.1439x^2^ - 11.892x + 32.767 (R2 = 0.6507). **(C)** Data of A normalized to the number of residues in each conservation group. The discontinuous line corresponds to function y = -0.0012x^2^ + 0.0103x + 0.0047 (R2 = 0.7036). **(D)** Data of B normalized to the number of residues in each conservation group. The discontinuous line corresponds to function y = 0.0117x0.4096 (R2 = 0.2824). **(E-H)** Same as A-D but at the amino acid level. In this case the possible conservation windows covered 0% to 100%. The defining functions are E: y = -10.39ln(x) + 21.797 (R2 = 0.7694); F: y = 0.9015x^2^ – 9.5106x + 23.7 (R2 = 0.7085); G: y = -0.0023x^2^ + 0.0222x + 0.0011 (R2 = 0.5193); H: y = 0.0127ln(x) + 0.0178 (R2 = 0.1291). The position of each heterogeneity site is given in Table S3 (http://babia.cbm.uam.es/~lab121/SupplMatGarcia-Crespo).

### Correlation between quasispecies residue variation and mutations recorded in HCV from infected patients

Sites of variation in HCV replicating within individual patients may not segregate into the same conservation groups than the sites that mutate in cell culture. To explore this possibility, we examined the conservation category of the 197 amino acids (comprised between amino acids 124 and 320 of NS5B), which included 177 variable positions in HCV from a cohort of 220 HCV-infected patients [22] (Table S4 and S5 in http://babia.cbm.uam.es/~lab121/SupplMatGarcia-Crespo). [The amino acid level is the one for which we had developed the informatics processing algorithms for HCVs of infected patients [22,30]]. The results show again a spread among conservation groups (Fig. 6). Thus, the HCV amino acid conservation pattern deduced from the LANL alignment does not fit the conservation observed in mutant spectra evolving in a cohort of infected patients.

**Figure 6.**
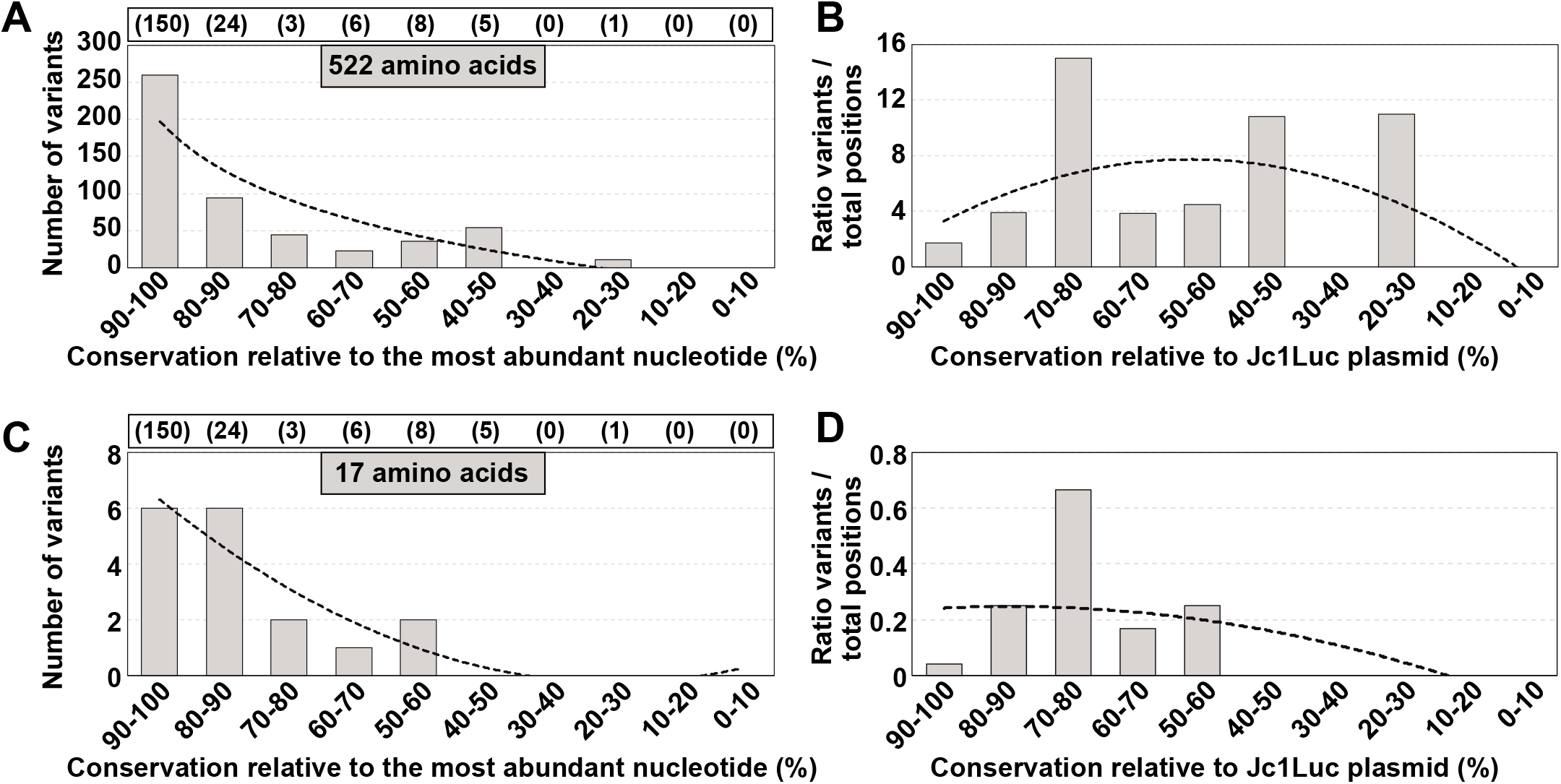
Distribution of positions with variant amino acids identified in HCV from infected patients among amino acid conservation groups according to the LANL amino acid sequence alignment. The residues under study span amino acid 124 to amino acid 320 of protein NS5B (which correspond to genomic nucleotides 7971 to 8561; H77 numbering). **(A)** Assignment of amino acid position where substitutions were located (listed in Table S5) (http://babia.cbm.uam.es/~lab121/SupplMatGarcia-Crespo) to amino acid conservation groups calculated from the amino acid sequence alignment of LANL. The 522 amino acids (within the 197 amino acids stretch) result from multiple substitutions at a given site due to the several HCV subtypes that infected the patients of the cohort under study [22] (shown also in Table S5 in http://babia.cbm.uam.es/~lab121/SupplMatGarcia-Crespo). Conservation groups are indicated in abscissa, and the number of variants is indicated in ordinate. The total number of amino acids that fall in each conservation category is indicated in parenthesis in the upper box. The discontinuous line corresponds to function y = -96.03ln(x) + 197.25 (R2 = 0.8028). **(B)** Same as A, but with values normalized to the total number of amino acids that falls within each conservation group. The defining function is y = - 0.3049x^2^ + 2.9414x + 0.6407 (R2 = 0.2448). **(C-D)** Same as A, B but with the 17 positions where a variant amino acid at a given position is common to infected patients and the HCV cell culture quasispecies. The defining functions are: C: y = 0.1326x^2^ – 2.1311x + 8.3167 (R2 = 0.8948); D: y = -0.0054x^2^ + 0.0219x + 0.2249 (R2 = 0.3209). In this case a calculation of conservation groups using as reference the HCV amino acid sequence encoded in plasmid Jc1Luc was not possible because the patients under study were not infected with HCV of the same subtype. See Table S5 (http://babia.cbm.uam.es/~lab121/SupplMatGarcia-Crespo) for the position of each amino acid substitution.

### Evidence of relaxed mutational acceptance in HCV quasispecies

One possibility to explain that nucleotides and amino acids that are conserved in consensus sequences or in the LANL alignment vary in mutant spectra is a higher tolerance for mutations during quasispecies dynamics than reflected in the consensus sequences or in data bank repositories. To address this point, we performed additional calculations, including conservation group resampling with some specific HCV genotypes, and simulations with randomly chosen genomic positions (Fig. S3 in http://babia.cbm.uam.es/~lab121/SupplMatGarcia-Crespo). We compared the conservation score of each position between residues 2769 and 9377 for selected HCV genotypes in the LANL alignment (H77 numbering) (Fig. S3A-D in http://babia.cbm.uam.es/~lab121/SupplMatGarcia-Crespo), and that of the heterogeneity sites. Both distributions were similar but with a larger accumulation of sites in the highest conservation window for the all-residue LANL calculation (compare Figs. 4A and S3A-H); the difference between them was statistically significant (p=0.0125, chi-square test with Monte Carlo correction; calculation made after reducing the 6669 residues proportionally to 114 to equate both data sets). The difference was also significant when the conservation groups were calculated using as reference the HCV sequence in Jc1Luc (p=0.0065, chi-square test with Monte Carlo correction; Fig. 4B and Fig. S3B in http://babia.cbm.uam.es/~lab121/SupplMatGarcia-Crespo). As a simulation, the distribution of conservation groups of 114 positions randomly chosen between residues 2769 and 9377 (H77 numbering) carried out in triplicate was also similar to the experimental distribution, but again with a significant accumulation of sites in the highest conservation window (Table S6, S7 and Fig. S3 I-K in http://babia.cbm.uam.es/~lab121/SupplMatGarcia-Crespo). Thus, the conservation groups of the variable sites in the HCV mutant spectra suggest a relaxed acceptance of mutations but significantly different from the prediction of a chance distribution of mutations.

### A possible alternative conservation criterion

The relaxed (but not unlimited) acceptance of mutations in evolving quasispecies hints at the possibility that a new criterion for conservation might result from sequence alignments in which all mutations found in mutant spectra are included, independently of their origin or frequency level. We have derived such alignment for NS5B amino acids 124 to 320, with inclusion of all available cell culture and *in vivo* sequences used in the present study. The results (Fig. 7) indicate that 5.1% of the amino acid positions are conserved in this new alignment. The proportion of conserved residues is dramatically reduced relative to the classical alignment with LANL sequences. Obviously, as additional mutant spectra of HCV populations are characterized by broad implementation of deep sequencing procedures in the clinic, the number of conserved residues is expected to diminish. Although absolute conservation cannot be foreseen as an attribute of the great majority of residues, the more stringent assessment of conservation attained with mutant spectrum alignments may provide a more realistic criterion on which to base the design of broad-spectrum antiviral ligands and vaccines.

**Figure 7.**
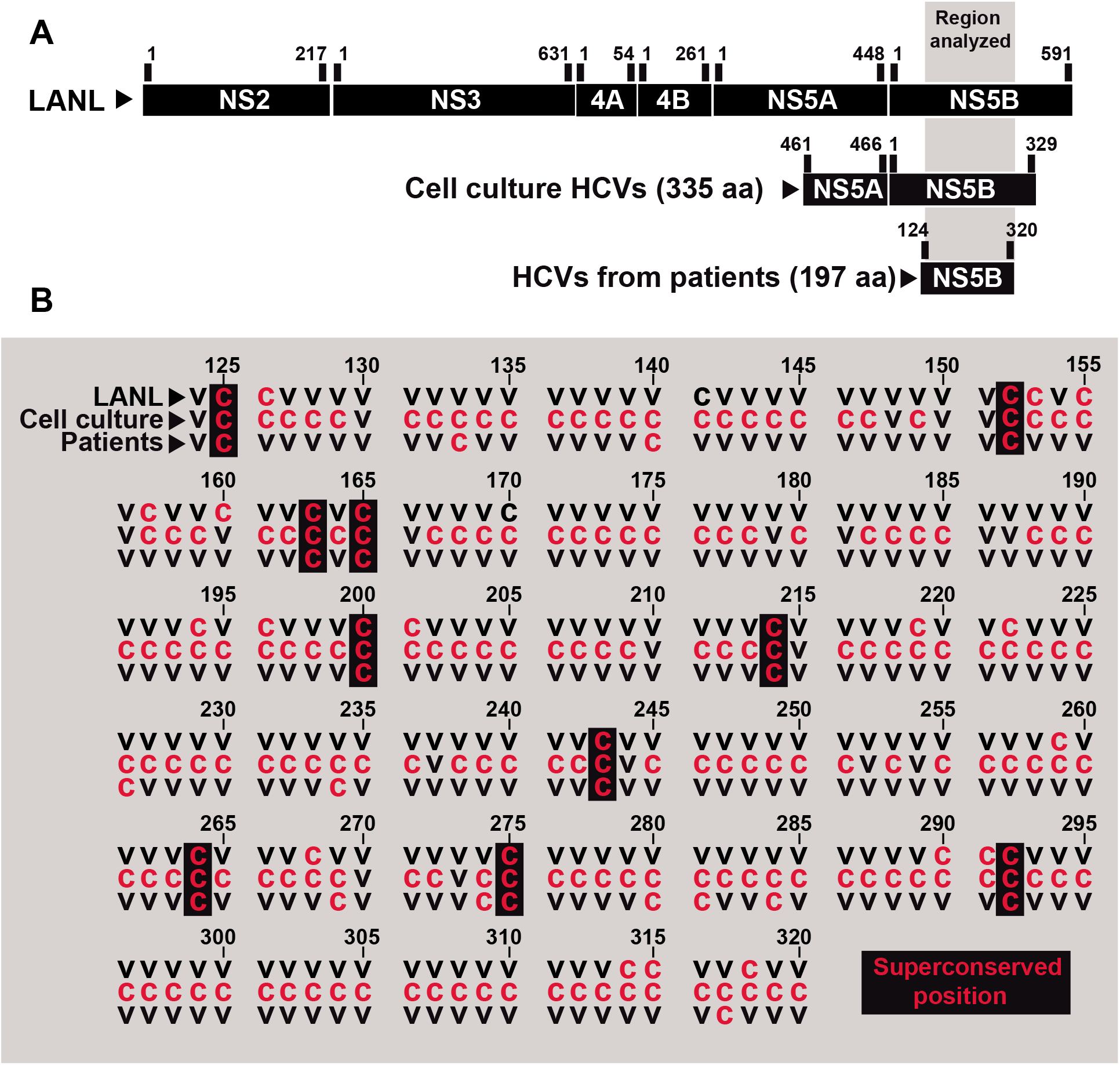
Distribution of superconserved positions among the amino acids 124 to 320 of NS5B (corresponding to genomic residues 7971 to 8561; numbering according to HCV reference isolate H77). **(A)** Scheme of the different regions and the number of amino acids analyzed from patients, cell culture or LANL. **(B)** Sequence alignment to indicate conserved positions. All amino acid substitutions identified in quasispecies of HCV evolving in cell culture and in infected patients, analyzed in the present study, have been considered to determine that a site is variable. Totally conserved positions are depicted by a red C, and variable positions (independently of the origin and extent of the variation) by a black V. Superconserved positions are indicated by a black rectangle.

## Discussion

Characterization of viral quasispecies *in vivo* is based mainly on intra-host mutant spectrum groupings of related sequences, time-dynamics analyses, and quantification of diversity indices [20,31–34]. These approaches have progressed quite independently from those employed to address long-term viral evolution, which are centered on phylogenetic and phylodynamic analyses of consensus sequences of independent viral isolates [6,7,35]. Attempts to connect the two domains of virus evolution have been limited [5]. The main objective of the present study has been to evaluate if conservation of HCV residues in consensus sequences or in data banks reflects also conservation at the level of mutant spectra of the populations. The comparisons have involved mutant spectra from a clonal HCV population passaged in cell culture, and viral populations from infected patients. In all cases examined residue conservation defined in consensus sequences was ampler than that in mutant spectra. A similar discordance was quantified when mutant spectra were compared with HCV sequences in the LANL data bank. This quantitatively important lack of correspondence between conservation scores is probably influenced by a relaxed negative selection acting on the newly arising mutations that do not reach dominance in the populations (frequencies below 50% in the mutant spectrum). In addition to the loss of resolution inherent to the conversion of a mutant spectrum into a consensus sequence, the great majority of sequences deposited in LANL belong to viruses that have undergone selective filters and bottleneck events in natural environments. Specifically, immune responses are evoked in most infected individuals and population bottlenecks accompany host-to-host transmissions, and probably also intra-host events [36]. A possible criticism to our observations is that the level of mutation analysis is so different between consensus sequences and mutant spectra that no matter which comparisons we do, the result will always be the same. This is not correct. In a recent study [37], we defined a subset of amino acid substitutions in HCV-infected patients, termed highly represented substitutions (HRS), identified in the same cohort of the present study [22]; in contrast with the complete set of substitutions, HRSs are distributed among intermediate conservation groups from the LANL alignment [37].

Differences in conservation between mutant spectra and consensus sequences or sequences in data banks emphasize two general points: (i) the relative (contingent) nature of residue conservation in viral genomes, and (ii) the simplification implied in the conversion of a mutant cloud (the real biological entity) into a single consensus sequence which is a weighted average of many different sequences. Concerning (i), in addition to the results reported here, there are other lines of evidence that support contingency of residue conservation or variation. For example, upon passage of HCV in Huh-7.5 cells, the NS5A- and NS5B-coding regions accumulated many mutations while the hypervariable regions 1 and 2 (HVR1 and HVR2) —defined as hypervariable on the basis of comparison of clinical isolates— remained largely invariant [17]. In this case, variation was probably conditional upon the virus having responded to the immune response in infected patients [17]. Another example was provided by foot-and-mouth disease virus (FMDV), which includes an Arg-Gly-Asp triplet exposed on its capsid that serves both as a receptor recognition and antigenic site. The triplet is highly conserved among natural isolates, and was considered essential; yet it varied and proved dispensable for infection in cell culture [38], and in some cases also for infection in vivo [39]. Concerning (ii), we have previously underlined that exclusion of mutant spectrum information in data banks represents a limitation for many biological studies with viruses that exhibit quasispecies dynamics [36,40]. Such limitation has also been expressed on theoretical grounds when reducing information conveyed by individual sequences into a consensus sequence that may not even exist in the population it intends to represent [41].

As a practical consequence of our conclusions, long-term residue constancy, as calculated from sequence alignments in data banks, does not guarantee a limitation in selection of escape mutants during intra-host evolution. It may be considered that some residues (i.e. those that are part of the catalytic site of viral enzymes) should be perfect ligands for inhibitors, no matter what the type of conservation correlations here described might indicate. However, ligand design is often based on several residues (or on structural elements that depend on several residues), and not all of them necessarily belong to a catalytic site; in addition, alteration of the conformation of an RNA structure or a protein catalytic domain by distant residues has been reported [42,43].

When a ligand is directed towards a viral population harboring a wide variant repertoire, selection and survival of ligand-escape mutants will be critically dependent on the viral population size and the number of different sequences endowed with comparable fitness level. The broad sequence repertoire may guide minority subpopulations towards dominance despite the transiency of the initial mutations that mediated the escape. Therefore, residue conservation in data banks should not endorse the design of viral ligands intended to control viral quasispecies. A new, more stringent criterion for conservation based on meta-quasispecies analyses with minority mutant spectra included in the alignments would seem more adequate. Even with this new tool, the random nature of mutations, their rate of occurrence, the positive and negative intra-genome interactions among mutations (epistasis), and inter genome interactions (complementation, cooperation, interference) render evolutionary pathways unpredictable and changeable [36,44]. Design of universal vaccines and antiviral agents (with potential of broad coverage to confront viral infections) is likely to remain an important challenge.

The elevated mutant repertoire that participates in quasispecies dynamics and its concealment in consensus sequences, together with evolutionary constraints at the epidemiological levels, may contribute to lower rates calculated for inter-host than intra-host evolution, one of the open questions in virus evolution [6,23–27]. Explorations of sequence space and opportunities for rapid short-term RNA virus evolution are far more abundant at the quasispecies level than reflected in data banks.

## Materials and Methods

### Experimental data on HCV quasispecies

The nucleotides in the HCV genome that participate in mutational waves or that belong to sites with composition heterogeneity in mutant spectra of virus evolving in cell culture (and deduced amino acid replacements) have been described [19]. For deep sequencing (Miseq platform, 2x300bp mode, v3 chemistry) basal sequencing errors, absence of PCR-mediated recombination, and haplotype frequency reproducibility were controlled experimentally and bioinformatically. Experimental controls involved deep sequencing of reconstructed mixtures of HCV RNAs containing different proportions of RNAs with specific mutations, comparison of results of mutant spectrum composition of the same viral sample subjected to two different amplifications and data processing cycles, and quantification of basal recombination rates upon amplification of marked RNAs [19,20,30]. Bioinformatically, controls were based on computations of binomial distributions for different coverages in the range of 500-10,000 reads [30]. The cut-off frequency of mutant detection was 0.5% [19].

HCV sequences from infected patients are those described in [22]. Amino acid substitutions belong to HCV from patients that failed direct acting antiviral agents, and no distinction has been made among mutant frequency values in the populations analyzed. Deep sequencing procedures and bioinformatics processing of data were the same applied to the HCV quasispecies in cell culture [22]. For the patient samples, the cut-off frequency of mutant detection was 1% [22,30]. Only mutations identified in the two DNA strands were included in the calculations.

Sites of nucleotide heterogeneity within the NS2-NS5B-coding region are those described in [19], without distinction of the frequency of the mutant nucleotide. Oligonucleotide primers, RT-PCR amplification conditions, and nucleotide sequencing (23 ABI 3730 XLS sequencer, Macrogen Inc.) were previously described [19].

### Mutation level and nomenclature

Individual mutations that change in frequency (those that participate in mutational waves) have been previously divided into three levels according to the frequency that they reached in the HCV population evolving in Huh-7.5 cells. Level L_0_ include those mutations that at some passage reached a frequency above the 0.5% cut-off level and that never exceeded 1% frequency in the sample. Level L1 corresponds to mutations whose frequency was never above 10%. Level L2 is the one of mutations that at some virus passage attained a frequency higher than 10%. The level assignment of each mutation was previously described [19], but no distinction among levels has been made for the present study.

Some of the mutations identified were present in more than one population [for example in populations from replicas (a), (b) or (c)] in cell culture which would qualify them as single nucleotide polymorphisms (SNPs) in general genetics nomenclature, while other mutations are present in only one lineage. The latter mutations deviate from the bona fide genetic polymorphism concept, and yet they contribute to the mutant swarm nature to the population. For the calculations performed in the present study no distinction has been made among mutations according to their L level or for being present in one or more lineages, and in consequence the term SNP is not used.

In addition to mutations, heterogeneous sites refer to those nucleotide positions in the consensus sequence determined by Sanger sequencing of a population in which two nucleotides were present. The criteria to establish a position as heterogeneous is that the two coexisting nucleotides are detected upon sequencing of the two cDNA strands, and that the mutant nucleotide exceeds a detection limit value of 15% [further details in [19]]. For the calculations of the present study no distinction has been made among heterogeneous sites according to the percentage of mutant nucleotide.

### Sequences from the Los Alamos data base

The sequences were retrieved from LANL following previously described procedures [22,30]. Inclusion criteria were that the sequences had been confirmed, that they corresponded to full-length (or near-full length) genomes (without large insertions or deletions), and with no evidence of their being recombinants. Their HCV genotype / subtype distribution is: 553 sequences of genotype G1a; 427 of G1b; 3 of G1c; 33 of G2a; 81 of G2b; 8 of G2c; 5 of G2j; 4 of G2k; 49 of G3a; 17 of G4a; 5 of G4d; and 6 of G4f. They were aligned using the program BioEdit version 7.0.9.0. For the calculation of the conservation range of individual residues, no distinction has been made between HCV subtypes. As part of the controls to exclude a bias in the assignment of variable sites to the different conservation groups, calculations were made with conservation groups determined with an alignment of sequences that included an equal representation of different subtypes.

### Statistics

The statistical significance of differences in the distribution of variable sites among conservation groups was calculated with the Pearson’s chi-square test using software R version 3.6.2, without or with Monte Carlo correction (based on 2000 replicates). Sample sizes are given for each comparison.

### Sequence accession numbers and data availability

The reference accession numbers of sequences retrieved from LANL used to determine conservation groups are given in Table S2 (http://babia.cbm.uam.es/~lab121/SupplMatGarcia-Crespo). Accession numbers for HCV samples included in the patient cohort are SAMN08741670 to SAMN08741673 [30]. Amino acid replacements in HCV from infected patients have been previously described [22], and are compiled in Table S5 in http://babia.cbm.uam.es/~lab121/SupplMatGarcia-Crespo. GenBank accession numbers for HCV p0, HCV p100 and HCV p200 are KC595606, KC595609 and KY123743, respectively. Illumina data can be retrieved from the NCBI BioSample data base, with accession numbers SAMN13531332 to SAMN13531367 (Bio Project accession number PRJNA593382).

## Acknowledgments

We are indebted to J.C. de la Torre and Matthias Pauthner for the critical reading of the manuscript and valuable comments. The work at CBMSO was supported by grants SAF2014-52400-R from Ministerio de Economía y Competitividad (MINECO), SAF2017-87846-R, BFU2017-91384-EXP from Ministerio de Ciencia, Innovación y Universidades (MCIU), PI18/00210 from Instituto de Salud Carlos III, S2013/ABI-2906, (PLATESA from Comunidad de Madrid/FEDER) and S2018/BAA-4370 (PLATESA2 from Comunidad de Madrid/FEDER). C.P. is supported by the Miguel Servet program of the Instituto de Salud Carlos III (CP14/00121 and CPII19/00001) cofinanced by the European Regional Development Fund (ERDF). CIBERehd (Centro de Investigación en Red de Enfermedades Hepáticas y Digestivas) is funded by Instituto de Salud Carlos III. Institutional grants from the Fundación Ramón Areces and Banco Santander to the CBMSO are also acknowledged. The team at CBMSO belongs to the Global Virus Network (GVN). The work in Barcelona was supported by Instituto de Salud Carlos III, cofinanced by the European Regional Development Fund (ERDF) grant number PI19/00301 and by the Centro para el Desarrollo Tecnológico Industrial (CDTI) from the MCIU, grant number IDI-20151125. Work at CAB was supported by MINECO grant BIO2016-79618R and PID2019-104903RB-I00 (funded by EU under the FEDER program) and by the Spanish State research agency (AEI) through Project number MDM-2017-0737 (Unidad de Excelencia “María de Maeztu”-Centro de Astrobiología (CSIC-INTA). C. G.-C. is supported by predoctoral contract PRE2018-083422 from MCIU. B. M.-G. is supported by predoctoral contract PFIS FI19/00119 from Instituto de Salud Carlos III (Ministerio de Sanidad y Consumo) cofinanced by Fondo Social Europeo (FSE).

