## Supplemental Material for "Dissimilar conservation pattern in hepatitis C virus mutant spectra, consensus sequences, and data banks"

### <u>Supplementary material for:</u>

**Table S1. Mutations and corresponding amino acid substitutions of the NS5A-NS5B-coding region in the mutant spectra of viral populations analyzed by ultra deep sequencing[a].**

| Mutation[b] | Amino Acid substitution[c] |
|---|---|
| A7650G[d] | **D461G** |
| U7651C[d] | Syn |
| U7651G[d] | **D461E** |
| A7652G[d] | **T462A** |
| A7652G[d]<br>C7653G[d]<br>C7654A[d] | **T462G** |
| A7652C[d]<br>C7653G[d]<br>C7954G[d] | **T462R** |
| A7655G[d] | **T463A** |
| A7655U[d] | **T463S** |
| C7656A[d] | **T463N** |
| C7656G[d] | **T463S** |
| G7658A[d] | **V464M** |
| G7658C[d] | **V464L** |
| G7658U[d] | **V464L** |
| U7659A[d] | **V464E** |
| U7661A[d] | **C465S** |
| U7661C[d] | **C465R** |
| C7681U | Syn |
| U7693G | Syn |
| A7696G | Syn |
| A7717G | Syn |
| U7727C | Syn |
| U7741C | Syn |
| G7744A | Syn |
| A7762C | Syn |
| U7768C | Syn |
| U7783C | Syn |
| A7786G | Syn |
| A7789G | Syn |
| U7802C | **S46P** |
| G7810A | Syn |
| U7825A | Syn |
| U7825C | Syn |
| U7842C | **V59A** |
| U7858C | Syn |
| A7864G | Syn |
| U7868C | Syn |
| A7870G | Syn |
| C7876U | Syn |
| G7886A | **A74T** |
| C7900G | Syn |

| | |
|---|---|
| C7903A | **S79R** |
| A7906C | Syn |
| A7906G | Syn |
| A7907U<br>G7908U | **R81L** |
| U7911G | **L82R** |
| C7915G | Syn |
| C7918U | Syn |
| U7919C | Syn |
| G7924A | Syn |
| G7930U | Syn |
| U7937C | Syn |
| U7942C | Syn |
| U7951C | Syn |
| U7954A | Syn |
| U7954C | Syn |
| A7957G | Syn |
| A7958G | **R98G** |
| A7965G | **K100R** |
| U7969C | Syn |
| A7972G | Syn |
| C7975U | Syn |
| G7978A | Syn |
| U7997C | Syn |
| U8000G | **S112A** |
| G8007A | **R114K** |
| C8020U | Syn |
| A8036G | **K124E** |
| U8068C | Syn |
| C8071U | Syn |
| A8074G | Syn |
| A8089G | Syn |
| U8092C | Syn |
| C8101U | Syn |
| G8107C | Syn |
| G8108A | **D148N** |
| G8114A | **A150T** |
| A8118G | **K151R** |
| U8125C | Syn |
| C8132G | **P156A** |
| C8132U | **P156S** |
| U8137C | Syn |
| A8144C | **I160L** |
| C8161U | Syn |
| G8162A | **G166S** |
| C8191U | Syn |
| U8200C | Syn |
| A8201G | **T179A** |
| G8209A | Syn |
| G8209U | **K181N** |
| U8215C | Syn |
| G8222A | **V186I** |
| U8233C | Syn |

| | |
|---|---|
| U8239C | Syn |
| C8242U | Syn |
| C8251U | Syn |
| C8260U | Syn |
| A8263G | Syn |
| U8275C | Syn |
| C8278G | Syn |
| C8278U | Syn |
| A8295G | **E210G** |
| U8310C | **M215T** |
| U8314G | Syn |
| U8317C | Syn |
| U8323C | Syn |
| U8326C | Syn |
| U8353C | Syn |
| G8374A | Syn |
| A8376G | **E237G** |
| C8386U | Syn |
| U8396C | **S244P** |
| C8398U | Syn |
| C8399U | Syn |
| C8404G | Syn |
| U8419C | Syn |
| C8421U | **A252V** |
| C8421U<br>C8422U | **A252V** |
| C8422U | Syn |
| A8427G | **H254R** |
| U8446G | Syn |
| A8455G | Syn |
| C8470U | Syn |
| A8475G | **K270R** |
| U8479C | Syn |
| A8483G | **T273A** |
| U8491C | Syn |
| U8491G | Syn |
| C8494U | Syn |
| G8518A | Syn |
| A8521U | Syn |
| C8530U | Syn |
| U8536C | Syn |
| C8539U | Syn |
| C8551U | Syn |
| A8566G | Syn |
| C8575U | Syn |
| A8577G | **K304R** |
| U8581C | Syn |
| G8584A | Syn |
| C8595A | **A310E** |
| A8602G | Syn |
| U8620C | Syn |
| A8626G | Syn |

| | |
|---|---|
| **Total Mutations[e]** | **145** |
| **Synonymous[f]** | **101** |
| **Non-synonymous[g]** | **44** |

[a] The HCV populations analysed and the location of the mutations, encoded amino acid substitutions and their tolerability were previously described in Gallego et al, J Virol 94(6), 2020 doi: 10.1128/JVI.01856-19.

[b] The HCV genome residue numbering corresponds to the JFH-1 genome (accession number #AB047639); genomic residues 7649 to 8653 were analysed.

[c] Amino acid residues (single letter code) are numbered for the C-terminal part of NS5A and from the N- to the C-terminus of NS5B.

[d] Mutations (and deduced amino acid substitutions) of NS5A.

[e] Number of different mutations found counted relative to the sequence of the parental plasmid Jc1FLAG2(p7-nsGluc2A).

[f] Number of different synonymous substitutions found counted relative to the sequence of the parental plasmid Jc1FLAG2(p7-nsGluc2A).

[g] Number of different non-synonymous substitutions found counted relative to the sequence of the parental plasmid Jc1FLAG2(p7-nsGluc2A).

**Table S2. Reference accession numbers of sequences retrieved from Los Alamos database.**

| Genotype 1 | |
|---|---|
| **Subtype 1a** | AB520610[a], AF009606[a], AF011751 - AF011753[a], AF271632[a], AF511949[c], AF511950[c], AJ278830, EF407411 - EF407415, EF407417 - EF407419, EF407421 - EF407423, EF407425, EF407427, EF407428, EF407431 - EF407447, EF407449 - EF407457, EF621489, EU155214 - EU155216, EU155233, EU155236 - EU155245, EU155249 - EU155252, EU155265 - EU155278, EU155282 - EU155288, EU155291, EU155293, EU155294, EU155296, EU155297, EU155299, EU155309, EU155311, EU155313, EU155314, EU155319 - EU155323, EU155338 - EU155355, EU155378, EU155379, EU234064, EU234065, EU239715, EU239716, EU250017, EU255927 - EU255958, EU255963 - EU255971, EU255973 - EU255992, EU255994 - EU255999, EU256002 - EU256024, EU256026, EU256028 - EU256034, EU256036 - EU256044, EU256047 - EU256053, EU256055 - EU256058, EU256060, EU256067, EU256068, EU256070 - EU256074, EU256087, EU256094, EU256095, EU256097, EU256105 - EU256107, EU260396, EU362876, EU362877, EU362879, EU362880, EU362882, EU362884 - EU362887, EU362891 - EU362898, EU362901, EU482831, EU482832, EU482834 - EU482838, EU482840 - EU482848, EU482852 - EU482858, EU482861 - EU482873, EU482878, EU482882, EU482884, EU482887, EU482889, EU529676 - EU529681, EU569722, EU569723, EU595697 - EU595699, EU660383 - EU660385, EU660387, EU687193 - EU687195, EU781746 - EU781803, EU781805 - EU781822, EU862824, EU862827, EU862828, EU862830 - EU862832, EU862834, EU862839 - EU862841, FJ024087, FJ024274 - FJ024276, FJ024278, FJ024280 - FJ024282, FJ181999 - FJ182001, FJ205867 - FJ205869, FJ390394, FJ390395, FJ390399, FJ410172, GQ149768, JQ914271, JQ914272, JX463525 - JX463530, JX463532 - JX463538, JX463541 - JX463545, JX463551 - JX463615, JX463617 - JX463622, JX463624 - JX463626, JX463628 - JX463633, JX463635 - JX463638, KC844049, M62321[a], M67463, NC_004102[a]. |
| **Subtype 1b** | AB049087 - AB049096, AB049098 - AB049101, AB080299[a], AB154177 - AB154206, AB191333, AB249644, AB426117[a], AB429050, AB435162[a], AB442219 - AB442222, AB691953, AB779562, AB779679, AF054247 - AF054259[a], AF165045 - AF165064, AF176573, AF207752 - AF207758, AF207760 - AF207774, AF208024, AF313916, AF333324[b], AF356827, AF483269, AJ000009, AJ132996[c], AJ132997, AJ238799, AJ238800, AY045702, AY587016[a], AY587844, D10750[b], D10934, D11168, D11355, D13558[b], D14484, D30613, D45172[a], D50480 - D50485, D63857, D85516, D89815, D89872[c], D90208, DQ071885, EF032892 - EF032894, EF407458 - EF407504, EU155217 - EU155232, EU155235, EU155253 - EU155264, EU155279 - EU155281, EU155300 - EU155308, EU155315 - EU155318, EU155324 - EU155337, EU155356 - EU155377, EU155381, EU155382, EU234061, |

| | |
|---|---|
| | EU234062, EU239714, EU255960 - EU255962, EU256000, EU256001, EU256045, EU256059, EU256061, EU256062, EU256064 - EU256066, EU256075 - EU256085, EU256088 - EU256092, EU256098 - EU256103, EU482833, EU482839, EU482849, EU482859, EU482860, EU482874, EU482875, EU482877, EU482879 - EU482881, EU482883, EU482885, EU482886, EU482888, EU529682, EU660386, EU660388, EU781825 - EU781832, EU862835, EU862837, FJ024086, FJ024277, FJ024279, FJ390396 - FJ390398, FJ478453, FN435993, GU133617 - GU451224, HQ110091, HQ639937, HQ639940, HQ639946, HQ639947, HQ719473, HQ912956 - HQ912959, JN120912, KC439481 - KC439527, KC844051, KC844052, L02836[c], M58335, M84754, M96362, U01214, U16362, U45476, X61596 |
| **Subtype 1c** | AY051292[c], D14853, KC844047 |
| **Genotype 2** | |
| **Subtype 2a** | AB047639 - AB047645, AB690460, AB690461[a], AF169002 - AF169005, AF177036[a], AF238481 - AF238485[c], AY746460[c], D00944[a], HQ639938, HQ639939, HQ639943 - HQ639945, JX014307[a], KC844043, KC967476, KF676351, KF676352, KF700370[a], NC_009823[c] |
| **Subtype 2b** | AB030907[a], AB559564, AB661373, AB661374, AB661376 - AB661378, AB661380 - AB661386, AB661389 - AB661393, AB661395 - AB661397, AB661399 - AB661403, AB661405 - AB661407, AB661409 - AB661422, AB661424 - AB661431, AF238486[c], AY232730 - AY232749, D10988, DQ430815, DQ430817, JQ745651[a], KC197226, KC844048, KC967477, KC967478 |
| **Subtype 2c** | D50409, JX227950, JX227951, JX227965, JX227966, KC197227, KC197228, KC967479 |
| **Subtype 2j** | HM777358, HM777359, JF735113, KC197232, KC197233 |
| **Subtype 2k** | AB031663, JX227952, JX227953, KC197234 |
| **Genotype 3** | |
| **Subtype 3a** | AB691595[a], AB691596[a], AB792683[a], AF046866[c], AY956467, D17763, D28917, DQ430819, DQ430820, DQ437509, GQ275355, GQ356200 - GQ356217, GU814263[c], HQ639941, HQ639942, HQ912953, JN714194, JQ717254 - JQ717260, KC844041, KF035123 - KF035127, NC_009824, X76918 |
| **Genotype 4** | |
| **Subtype 4a** | AB795432, DQ418782 - DQ418784, DQ418787 - DQ418789, DQ988073 - DQ988079, GU814265[c], NC_009825, Y11604 |
| **Subtype 4d** | DQ418786, DQ516083, EU392172, FJ462437, KC844045 |
| **Subtype 4f** | EF589160, EF589161, EU392169, EU392170, EU392174, EU392175 |

| | |
|---|---|
| **Infected patients** | **1136 (95.4%)** |
| **Clones** | **35 (2.9%)** |
| **Infected chimpanzees** | **3 (0.3%)** |
| **Undefined origin** | **17 (1.4%)** |
| **Total sequences** | **1191** |

[a] Sequences corresponding to clones.

[b] Sequences corresponding to infected chimpanzee.

[c] Sequences of undefined origin.

**Table S3. Mutations and corresponding amino acid substitutions deduced from the genomic sites with composition heterogeneity of the beginning of the NS2-coding region to the end of the NS5B-coding region in the genomic consensus sequence analysed by Sanger sequencing[a].**

| Mutation[b] | Amino Acid substitution[c] |
|---|---|
| U2838U/G | **F20F/C** |
| A2861A/G | **T28T/A** |
| U3001C/U | Syn |
| G3119A/G | **A114T/A** |
| C3271C/U | Syn |
| C3280C/U | Syn |
| U3550U/G | Syn |
| A3565A/U | Syn |
| U3674C/U | Syn |
| G3724A/G | Syn |
| A3802A/G | Syn |
| G3870A/G | **R147K/R** |
| C4099U/C | Syn |
| G4111G/A | Syn |
| C4159C/G | Syn |
| A4195G/A | Syn |
| G4216G/A | Syn |
| C4264G/C | Syn |
| A4286G/A | **I286V/I** |
| A4381G/A | Syn |
| A4402A/G | Syn |
| G4458A/G | **R343Q/R** |
| A4545A/C | **K372K/T** |
| U4591U/C | Syn |
| U4864U/C | Syn |
| G4954G/C | **E508E/D** |
| A4972A/G | Syn |
| U5200C/U | Syn |
| C5230U/C | Syn |
| G5378G/A | **A19A/T** |
| U5392U/C | Syn |
| A5396G/A | **I25V/I** |
| U5500U/C | Syn |
| C5506C/U | Syn |
| U5534U/C | Syn |
| A5551A/U | **Q22Q/H** |
| A5680G/A | Syn |
| A5683G/A | Syn |
| U5848U/C | Syn |
| A5954G/A | **I157V/I** |

| | |
|---|---|
| G6031G/A | Syn |
| G6126G/A | **R214R/K** |
| A6338G/A | **T24A/T** |
| U6350U/A | **F28F/I** |
| C6376U/C | Syn |
| C6412C/A | **A48A/D** |
| G6484G/U | **M72M/I** |
| A6491G/A | **T75A/T** |
| U6532U/C | Syn |
| A6636A/G | **Q123Q/R** |
| A6658A/G | Syn |
| A6686A/C | **I140I/L** |
| A6711A/U | **E148E/V** |
| G6732G/U | **G155G/V** |
| U6733U/C | Syn |
| G6748A/G | Syn |
| C6968A/C | **L234I/L** |
| A7001A/G | **T245T/A** |
| C7009G/C | **S247R/S** |
| G7081U/G | **E271D/E** |
| C7083C/A | **S272S/Y** |
| U7107U/C | **L280L/P** |
| A7110A/C | **E281E/A** |
| A7175G/A | **S303G/S** |
| U7181U/C | **F305F/L** |
| A7207A/U | Syn |
| A7218G/A | **Y317C/Y** |
| U7238A/U | **S324T/S** |
| A7302A/G | **K345K/R** |
| A7325G/A | **R353G/R** |
| G7345G/C | Syn |
| G7425A/G | **G386D/G** |
| C7444C/U | Syn |
| C7444C/A | Syn |
| A7452A/G | **E395E/G** |
| G7475G/A | **G403G/S** |
| A7494A/G | **E409E/G** |
| A7498A/G | Syn |
| G7499A/G | **G411S/G** |
| U7610U/C[d] | **S448S/P** |
| G7618G/A[d] | Syn |
| G7649G/A[e] | **D461D/N** |
| A7652G/A[e] | **T462A/T** |
| A7655U/A[e] | **T463S/T** |
| G7658A/G[e] | **V464M/V** |
| U7661A/U[e] | **C465S/C** |
| A7814A/G[e] | **K50K/E** |
| U7842U/C[e] | **V59V/A** |

| | |
|---|---|
| G7897G/A[e] | Syn |
| U7942C/U[e] | Syn |
| C7953C/G[e] | **S96S/C** |
| A7982A/G[e] | **K106K/E** |
| C8054C/G[e] | **P130P/A** |
| C8071C/U[e] | Syn |
| C8132U/C[e] | **P156S/P** |
| A8201A/G[e] | **T179T/A** |
| G8209U/G[e] | **K181N/K** |
| G8222G/A[e] | **V186V/I** |
| G8227G/A[e] | **M187M/I** |
| U8310C/U[e] | **M215T/M** |
| U8353C/U[e] | Syn |
| C8422U/C[e] | Syn |
| A8427A/G[e] | **H254H/R** |
| U8446G/U[e] | Syn |
| A8475A/G[e] | **K270K/R** |
| C8575C/U[e] | Syn |
| U8722A/U | **D352E/D** |
| A8752G/A | Syn |
| A8758G/A | Syn |
| C8854C/A | Syn |
| G8893G/A | Syn |
| U8929C/U | Syn |
| U9014G/U | **S450A/S** |
| C9385G/C | Syn |
| **Total Mutations[f]** | **114** |
| **Synonymous[g]** | **53** |
| **Non-synonymous[h]** | **61** |

[a] The HCV populations analysed, the location of the mutations, and examples of sites with two nucleotides were previously described in Gallego et al, J Virol 94(6), 2020 doi: 10.1128/JVI.01856-19.

[b] The HCV genome residue numbering corresponds to the JFH-1 genome (accession number #AB047639); genomic residues 2780 to 9442 were analysed.

[c] Amino acid residues (single letter code) are numbered from N- to the C-terminus of each region; Syn means synonymous (no amino acid replacement).

[d] Residues located in the insertion found in the NS5A region of genotype 2.

[e] Residues were mutational waves were also analysed.

[f] Number of different mutations found counted relative to the sequence of the parental plasmid Jc1FLAG2(p7-nsGluc2A).

[g] Number of different synonymous substitutions found counted relative to the sequence of the parental plasmid Jc1FLAG2(p7-nsGluc2A).

[h] Number of different non-synonymous substitutions found counted relative to the sequence of the parental plasmid Jc1FLAG2(p7-nsGluc2A).

**Table S4. Number of patients infected by each HCV subtype.**

| Genotype | Subtype | Number of patients |
|---|---|---|
| **G1** | **1a** | 50 |
| | **1b** | 87 |
| | **1l** | 1 |
| **G2** | **2c** | 1 |
| | **2i** | 2 |
| | **2j** | 1 |
| **G3** | **3a** | 47 |
| **G4** | **4d** | 27 |
| **Mixed infections** | **4d (69.3%) + 1b (30.6%)** | 1 |
| | **4a (98.5%) + 1b (1.5%)** | 1 |
| | **1b (80.1%) + 1a (19.9%)** | 1 |
| | **4d (96.7%) + 3a (3.3%)** | 1 |
| | **TOTAL** | **220** |

**Table S5. Location within NS5B amino acids 124 to 320 of the positions where amino acid substitutions were identified in infected patients[a].**

| Amino acid position[b] | Amino acid change | Number of variants[c] |
|---|---|---|
| **124[d]** | Yes | 10 |
| **125** | No | - |
| **126** | Yes | 2 |
| **127** | Yes | 3 |
| **128** | Yes | 4 |
| **129** | Yes | 2 |
| **130** | Yes | 7 |
| **131** | Yes | 9 |
| **132** | Yes | 1 |
| **133** | No | - |
| **134** | Yes | 2 |
| **135** | Yes | 5 |
| **136** | Yes | 2 |
| **137** | Yes | 1 |
| **138** | Yes | 1 |
| **139** | Yes | 2 |
| **140** | No | - |
| **141** | Yes | 1 |
| **142** | Yes | 3 |
| **143** | Yes | 3 |
| **144** | Yes | 1 |
| **145** | Yes | 3 |
| **146** | Yes | 3 |
| **147** | Yes | 4 |
| **148[d]** | Yes | 6 |
| **149** | Yes | 1 |
| **150[d]** | Yes | 7 |
| **151[d]** | Yes | 3 |
| **152** | No | - |
| **153** | Yes | 1 |
| **154** | Yes | 2 |
| **155** | Yes | 2 |
| **156[d]** | Yes | 3 |
| **157** | Yes | 1 |
| **158** | Yes | 1 |
| **159** | Yes | 1 |
| **160** | Yes | 2 |
| **161** | Yes | 1 |
| **162** | Yes | 4 |
| **163** | No | - |

| | | |
|---|---|---|
| **164** | Yes | 1 |
| **165** | No | - |
| **166[d]** | Yes | 1 |
| **167** | Yes | 3 |
| **168** | Yes | 1 |
| **169** | Yes | 4 |
| **170** | Yes | 1 |
| **171** | Yes | 7 |
| **172** | Yes | 2 |
| **173** | Yes | 4 |
| **174** | Yes | 2 |
| **175** | Yes | 2 |
| **176** | Yes | 2 |
| **177** | Yes | 4 |
| **178** | Yes | 4 |
| **179[d]** | Yes | 4 |
| **180** | Yes | 7 |
| **181[d]** | Yes | 6 |
| **182** | Yes | 3 |
| **183** | Yes | 2 |
| **184** | Yes | 11 |
| **185** | Yes | 4 |
| **186** | Yes | 3 |
| **187** | Yes | 3 |
| **188** | Yes | 2 |
| **189** | Yes | 10 |
| **190** | Yes | 4 |
| **191** | Yes | 1 |
| **192** | Yes | 1 |
| **193** | Yes | 2 |
| **194** | Yes | 1 |
| **195** | Yes | 2 |
| **196** | Yes | 3 |
| **197** | Yes | 1 |
| **198** | Yes | 5 |
| **199** | Yes | 1 |
| **200** | No | - |
| **201** | Yes | 1 |
| **202** | Yes | 3 |
| **203** | Yes | 2 |
| **204** | Yes | 1 |
| **205** | Yes | 3 |
| **206** | Yes | 10 |

| | | |
|---|---|---|
| **207** | Yes | 2 |
| **208** | Yes | 1 |
| **209** | Yes | 5 |
| **210[d]** | Yes | 8 |
| **211** | Yes | 2 |
| **212** | Yes | 3 |
| **213** | Yes | 8 |
| **214** | No | - |
| **215[d]** | Yes | 4 |
| **216** | Yes | 1 |
| **217** | Yes | 2 |
| **218** | Yes | 3 |
| **219** | Yes | 2 |
| **220** | Yes | 2 |
| **221** | Yes | 1 |
| **222** | Yes | 2 |
| **223** | Yes | 1 |
| **224** | Yes | 2 |
| **225** | Yes | 1 |
| **226** | No | - |
| **227** | Yes | 1 |
| **228** | Yes | 2 |
| **229** | Yes | 1 |
| **230** | Yes | 2 |
| **231** | Yes | 6 |
| **232** | Yes | 1 |
| **233** | Yes | 2 |
| **234** | No | - |
| **235** | Yes | 6 |
| **236** | Yes | 3 |
| **237[d]** | Yes | 1 |
| **238** | Yes | 3 |
| **239** | Yes | 3 |
| **240** | Yes | 2 |
| **241** | Yes | 1 |
| **242** | Yes | 2 |
| **243** | No | - |
| **244** | Yes | 5 |
| **245** | Yes | 4 |
| **246** | Yes | 5 |
| **247** | Yes | 2 |
| **248** | Yes | 3 |
| **249** | Yes | 2 |
| **250** | Yes | 3 |

| | | |
|---|---|---|
| **251** | Yes | 5 |
| **252[d]** | Yes | 4 |
| **253** | Yes | 3 |
| **254[d]** | Yes | 7 |
| **255** | Yes | 3 |
| **256** | Yes | 1 |
| **257** | Yes | 2 |
| **258** | Yes | 4 |
| **259** | Yes | 2 |
| **260** | Yes | 1 |
| **261** | Yes | 1 |
| **262** | Yes | 4 |
| **263** | Yes | 1 |
| **264** | No | - |
| **265** | Yes | 1 |
| **266** | Yes | 2 |
| **267** | Yes | 5 |
| **268** | Yes | 1 |
| **269** | No | - |
| **270[d]** | Yes | 4 |
| **271** | Yes | 2 |
| **272** | Yes | 4 |
| **273** | Yes | 5 |
| **274** | No | - |
| **275** | No | - |
| **276** | Yes | 4 |
| **277** | Yes | 1 |
| **278** | Yes | 3 |
| **279** | Yes | 2 |
| **280** | No | - |
| **281** | No | - |
| **282** | Yes | 2 |
| **283** | Yes | 1 |
| **284** | No | - |
| **285** | Yes | 5 |
| **286** | Yes | 3 |
| **287** | Yes | 1 |
| **288** | Yes | 2 |
| **289** | Yes | 2 |
| **290** | Yes | 1 |
| **291** | Yes | 2 |
| **292** | No | - |
| **293** | Yes | 3 |

| | | |
|---|---|---|
| **294** | Yes | 1 |
| **295** | Yes | 1 |
| **296** | Yes | 1 |
| **297** | Yes | 4 |
| **298** | Yes | 1 |
| **299** | Yes | 1 |
| **300** | Yes | 11 |
| **301** | Yes | 1 |
| **302** | Yes | 1 |
| **303** | Yes | 3 |
| **304[d]** | Yes | 4 |
| **305** | Yes | 2 |
| **306** | Yes | 2 |
| **307** | Yes | 7 |
| **308** | Yes | 2 |
| **309** | Yes | 4 |
| **310[d]** | Yes | 6 |
| **311** | Yes | 5 |
| **312** | Yes | 6 |
| **313** | Yes | 5 |
| **314** | Yes | 3 |
| **315** | Yes | 1 |
| **316** | Yes | 4 |
| **317** | No | - |
| **318** | Yes | 2 |
| **319** | Yes | 1 |
| **320** | Yes | 1 |
| **Total of positions with amino acid changes** | | **177** |
| **Total of positions without amino acid changes** | | **20** |
| **Total of variants** | | **522** |

[a] The clinical history of the 220 patients cohort under study was described in Chen et al., Antiviral Research 174: 104694, 2020.

[b] The HCV genome residue numbering corresponds to the H77 genome (accession number #AF009606); genomic residues 7971 to 8561 were analysed.

[c] Number of different amino acid changes detected in each position relative to the respective reference sequence. Multiple amino acid substitutions per site are explained by the different HCV genotypes and subtypes of the sequence under study (Chen et al., Antiviral Research 174: 104694, 2020).

[d] Position where a variant amino acid is common in an infected patients and some cell culture mutant spectrum.

**Table S6. Random distribution of positions in the HCV genome.**

| Randomized positions[a] | | |
|---|---|---|
| **Control 1[b]** | **Control 2[b]** | **Control 3[b]** |
| 2819 | 2799 | 2781 |
| 2906 | 2820 | 2840 |
| 2942 | 2866 | 2896 |
| 3093 | 2900 | 3048 |
| 3133 | 2904 | 3085 |
| 3220 | 2933 | 3407 |
| 3233 | 2961 | 3442 |
| 3272 | 3112 | 3603 |
| 3279 | 3146 | 3626 |
| 3360 | 3274 | 3628 |
| 3367 | 3361 | 3669 |
| 3453 | 3362 | 3690 |
| 3471 | 3399 | 3693 |
| 3493 | 3421 | 3694 |
| 3648 | 3467 | 3703 |
| 3657 | 3483 | 3733 |
| 3667 | 3705 | 3754 |
| 3669 | 3768 | 3837 |
| 3697 | 3839 | 3842 |
| 3882 | 3907 | 3916 |
| 4095 | 4034 | 3970 |
| 4103 | 4128 | 4008 |
| 4304 | 4133 | 4103 |
| 4332 | 4140 | 4149 |
| 4489 | 4158 | 4184 |
| 4517 | 4233 | 4190 |
| 4551 | 4303 | 4194 |
| 4761 | 4304 | 4202 |
| 4772 | 4305 | 4212 |
| 4812 | 4340 | 4455 |
| 4830 | 4362 | 4459 |
| 4998 | 4385 | 4471 |
| 5143 | 4397 | 4489 |
| 5155 | 4418 | 4568 |
| 5223 | 4432 | 4700 |
| 5246 | 4434 | 4784 |
| 5247 | 4537 | 4795 |
| 5296 | 4543 | 4918 |
| 5324 | 4551 | 4996 |
| 5393 | 4696 | 5058 |
| 5394 | 4711 | 5126 |

| | | |
|---|---|---|
| 5493 | 4747 | 5157 |
| 5519 | 4766 | 5264 |
| 5745 | 4814 | 5323 |
| 5750 | 5168 | 5414 |
| 5799 | 5288 | 5485 |
| 5912 | 5335 | 5492 |
| 5994 | 5456 | 5535 |
| 6051 | 5528 | 5610 |
| 6054 | 5551 | 5687 |
| 6220 | 5569 | 5851 |
| 6394 | 5572 | 5878 |
| 6512 | 5603 | 5913 |
| 6597 | 5695 | 6028 |
| 6630 | 5769 | 6078 |
| 6647 | 5821 | 6231 |
| 6693 | 5851 | 6256 |
| 6705 | 5937 | 6388 |
| 6764 | 6019 | 6427 |
| 6765 | 6046 | 6522 |
| 6777 | 6131 | 6542 |
| 6794 | 6149 | 6562 |
| 6826 | 6171 | 6593 |
| 6879 | 6173 | 6601 |
| 6890 | 6203 | 6616 |
| 6910 | 6244 | 6625 |
| 6934 | 6270 | 6776 |
| 6959 | 6282 | 6788 |
| 6969 | 6388 | 6803 |
| 7054 | 6389 | 6955 |
| 7201 | 6443 | 7024 |
| 7266 | 6771 | 7069 |
| 7267 | 6805 | 7085 |
| 7426 | 6818 | 7132 |
| 7438 | 6825 | 7145 |
| 7613 | 6837 | 7207 |
| 7677 | 6904 | 7247 |
| 7726 | 6923 | 7283 |
| 7806 | 6956 | 7569 |
| 7810 | 7025 | 7575 |
| 7948 | 7062 | 7638 |
| 7950 | 7076 | 7671 |
| 8003 | 7090 | 7783 |
| 8064 | 7131 | 7861 |
| 8109 | 7139 | 7897 |
| 8126 | 7145 | 7966 |
| 8148 | 7187 | 8049 |

| | | |
|---|---|---|
| 8156 | 7207 | 8060 |
| 8164 | 7286 | 8064 |
| 8195 | 7370 | 8131 |
| 8309 | 7397 | 8145 |
| 8357 | 7514 | 8385 |
| 8379 | 7577 | 8402 |
| 8390 | 7584 | 8420 |
| 8562 | 7733 | 8569 |
| 8622 | 7983 | 8602 |
| 8667 | 8002 | 8632 |
| 8690 | 8168 | 8675 |
| 8701 | 8180 | 8695 |
| 8711 | 8194 | 8709 |
| 8740 | 8304 | 8721 |
| 8750 | 8468 | 8736 |
| 8751 | 8505 | 8760 |
| 8789 | 8663 | 8772 |
| 8797 | 8707 | 8808 |
| 8815 | 8827 | 8881 |
| 8836 | 8830 | 8920 |
| 8868 | 8926 | 9038 |
| 8883 | 8927 | 9047 |
| 8939 | 8933 | 9178 |
| 9226 | 9021 | 9195 |
| 9233 | 9106 | 9279 |
| 9335 | 9117 | 9362 |
| 9337 | 9292 | 9378 |

[a]The HCV genome residue numbering corresponds to the JFH-1 genome (accession number #AB047639); genomic residues 2780 to 9442 were analysed.

[b] The assignments were as directed by Excel 2016.

**Table S7. Statistical analysis of the different distributions identified between HCV genomic residues 2769 (beginning of the NS2-coding region) and 9377 (end of the NS5B-coding region) (H77 numbering).**

| Comparison | Panel[a] | p-value<br>Chi-square test Monte Carlo correction | Significance[b] |
|---|---|---|---|
| Distribution of heterogeneity sites relative to most abundant nucleotide | 3A | 0.0165 | * |
| Random position distribution relative to the most abundant nucleotide | S2I | | |
| Distribution of heterogeneity sites relative to Jc1Luc plasmid | 3B | 0.0045 | ** |
| Random position distribution relative to Jc1Luc plasmid | S2J | | |
| Distribution of positions from Los Alamos alignment relative to the most abundant nucleotide | S2A | 0.9725 | ns |
| Random position distribution relative to the most abundant nucleotide | S2I | | |
| Distribution of positions from Los Alamos alignment relative to Jc1Luc plasmid | S2B | 1.0000 | ns |
| Random position distribution relative to Jc1Luc plasmid | S2J | | |

[a] Panel means figure number and panel in main text or supplemental material (S).

[b] The statistical significance of the differences is given as follows: ns, not significant; *, $P \leq 0.05$; **, $P < 0.01$.

### Figure S1

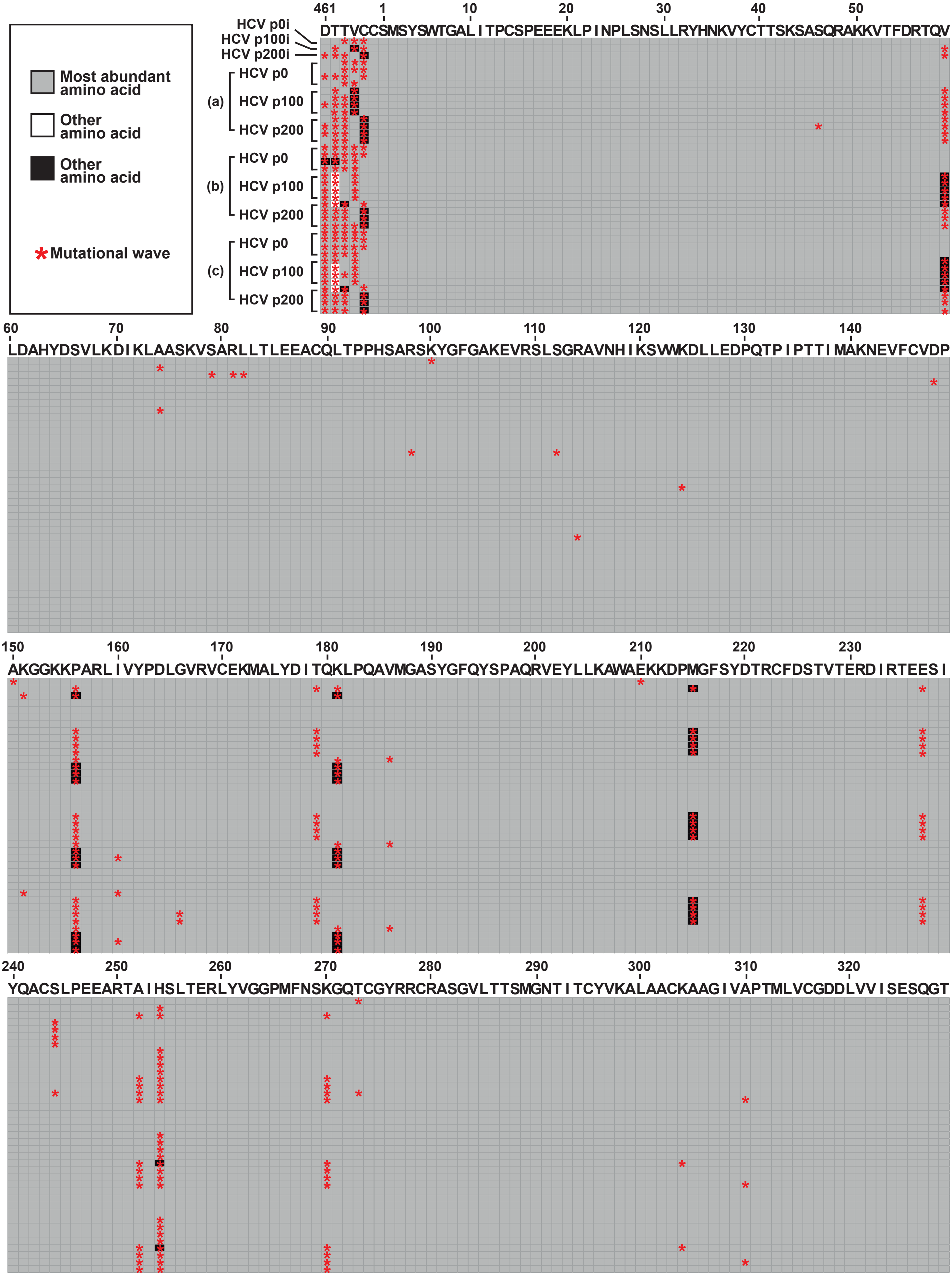
Most abundant amino acid
Other amino acid
Other amino acid
Mutational wave
HCV p0i
HCV p100i
HCV p200i
(a)
(b)
(c)
HCV p0
HCV p100
HCV p200
DTTVCCSMSYSWTGALITPCSPEEEKLPINPLSNSLLRYHNKVYCTTSKSASQRAKKVTFDRTQV
LDAHYDSVLKDIKLAASKVSARLLTLEEACQLTPPHSARSKYGFGAKEVRSLSGRAVNHIKSVWKDLLEDPQTPIPTTIMAKNEVFCVDP
AKGGKKPARLIVYPDLGVRVCEKMALYDITQKLPQAVMGASYGFQYSPAQRVEYLLKAWAEKKDPMGFSYDTRCFDSTVTERDIRTEESI
YQACSLPEEARTAIHSLTERLYVGGPMFNSKGQTCGYRRCRASGVLTTSMGNTITCYVKALAACKAAGIVAPTMLVCGDDLVVISESQGT

**Figure S1.** Heat map of an alignment of the 39 consensus amino acid sequences determined for the NS5A-NS5B region [encoded by nucleotides 7649 to 8653; numbering according to isolate JFH-1 (GenBank accession number #AB047639). The map for amino acids spans the same region for nucleotides shown in Fig. 2 of the main text. Each horizontal line represents the sequence of a population, as written on the left. The three first lines correspond to the initial populations, and the blocs below them include the populations at passages 1 to 4 of replicas (a), (b) and (c) of the three viral populations. Each column represents one of the 335 amino acid positions analyzed. Amino acid numbers for each protein are written above the amino acid sequence, which displays the most represented amino acid at each position in the sequences under comparison. The upper left box indicates the code for amino acid abundance: grey means an abundant amino acid (present in 66.7 % to 100 % of the compared sequences); white and black represent others amino acids presents in the same position. In NS5A a black square means the presence of amino acid E, R, S, M, and S at positions 461, 462, 463, 464, and 465, respectively, and white squares indicate the presence of amino acid A at position 462. In NS5B, black squares indicate the presence of amino acids A, S, N, T, and R, at positions 59, 156, 181, 215, and 254, respectively. Red asterisks point to amino acids encoded by nucleotides that participated in mutational waves (depicted in Fig. 2 of main text, and compiled in Table S1).

### Figure S2

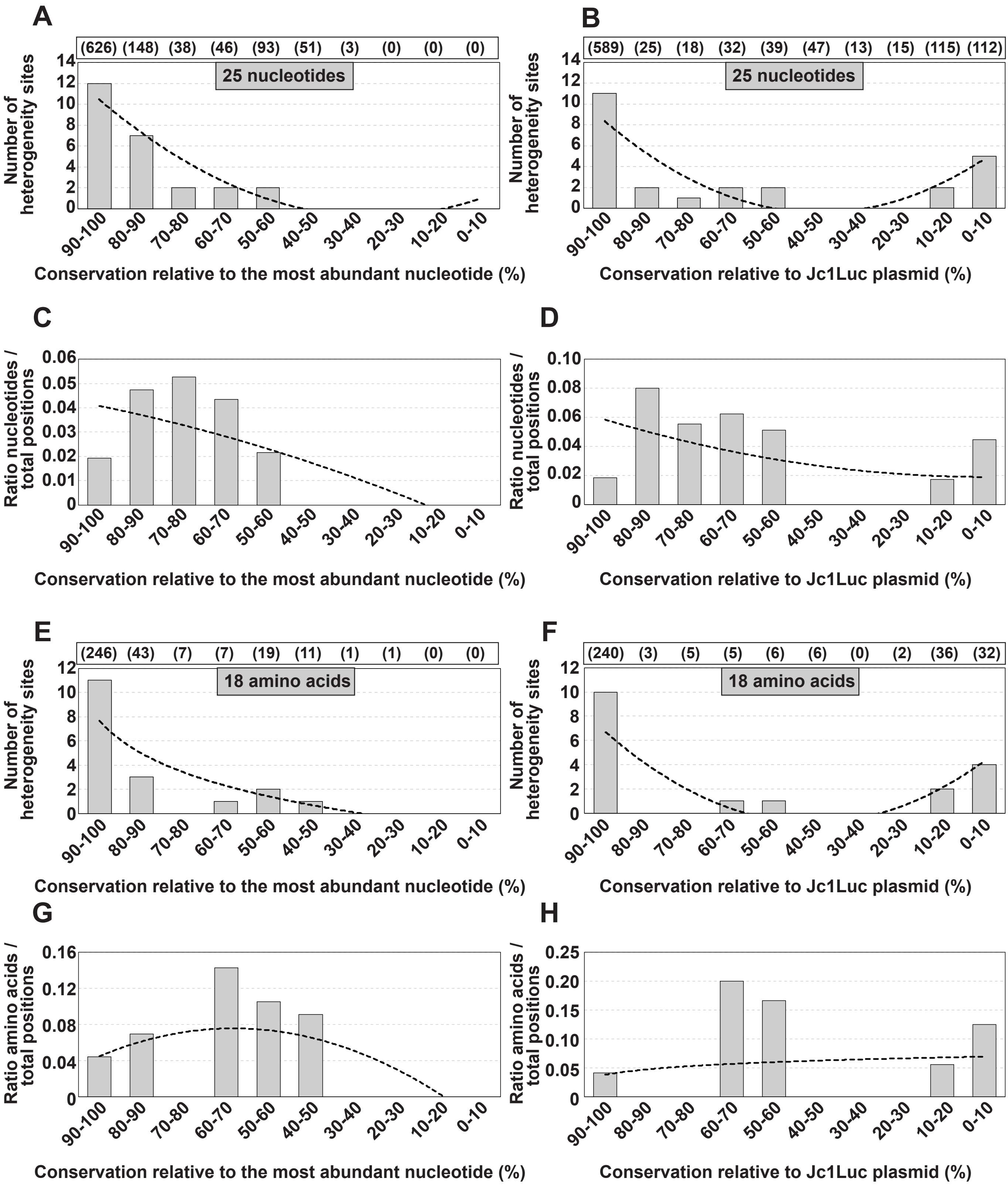
A
(626) (148) (38) (46) (93) (51) (3) (0) (0) (0)
25 nucleotides
Number of heterogeneity sites
90-100 80-90 70-80 60-70 50-60 40-50 30-40 20-30 10-20 0-10
Conservation relative to the most abundant nucleotide (%)
B
(589) (25) (18) (32) (39) (47) (13) (15) (115) (112)
25 nucleotides
Number of heterogeneity sites
Conservation relative to Jc1Luc plasmid (%)
C
Ratio nucleotides / total positions
Conservation relative to the most abundant nucleotide (%)
D
Ratio nucleotides / total positions
Conservation relative to Jc1Luc plasmid (%)
E
(246) (43) (7) (7) (19) (11) (1) (1) (0) (0)
18 amino acids
Number of heterogeneity sites
Conservation relative to the most abundant nucleotide (%)
F
(240) (3) (5) (5) (6) (6) (0) (2) (36) (32)
18 amino acids
Number of heterogeneity sites
Conservation relative to Jc1Luc plasmid (%)
G
Ratio amino acids / total positions
Conservation relative to the most abundant nucleotide (%)
H
Ratio amino acids / total positions
Conservation relative to Jc1Luc plasmid (%)

**Figure S2.** Number of nucleotides within HCV genomic residues 7584-8588 (H77 numbering) and deduced amino acid mixtures that belonged to heterogeneity sites, distributed among conservation groups according to the Los Alamos data base alignment. **(A)** Number of nucleotides at heterogeneity sites distributed among conservation groups, calculated relative to the most abundant nucleotide at the corresponding position in the Los Alamos alignment. The total number of nucleotides that fall in each conservation category is indicated in parenthesis in the upper box. The discontinuous line corresponds to function $y = 0.2652x^2 – 3.9773x + 14.167$ ($R^2 = 0.9117$). **(B)** Same as A but with nucleotide conservation in the Los Alamos alignment calculated relative to the corresponding residues in plasmid Jc1FLAG2(p7-nsGLuc2A (Marukian et al., Hepatology 48:1843-1850, 2008). The discontinuous line corresponds to function $y = 0.3371x^2 – 4.1144x + 12.15$ ($R^2 = 0.7324$). **(C)** Data of A normalized to the number of residues in each conservation group. The discontinuous line corresponds to function $y = -0.0002x^2 – 0.0031x + 0.0441$ ($R^2 = 0.5957$). **(D)** Data of B normalized to the number of residues in each conservation group. The discontinuous line corresponds to function $y = 0.0005x^2 – 0.0096x + 0.0676$ ($R^2 = 0.2183$). **(E-H)** Same as A-D but at the amino acid level. The defining functions are E: $y = -3.869\ln(x) + 7.6443$ ($R^2 = 0.6988$); F: $y = 0.3068x^2 – 3.6417x + 10.017$ ($R^2 = 0.6202$); G: $y = -0.0031x^2 + 0.0255x + 0.0227$ ($R^2 = 0.3955$); H: $y = 0.0136\ln(x) + 0.0383$ ($R^2 = 0.0168$). The position of each mutation and amino acid substitution deduced from the sites displaying composition heterogeneity is given in Table S3. The number of sequences retrieved from the Los Alamos data bank and inclusion criteria are explained in the main text.

**Figure S3**

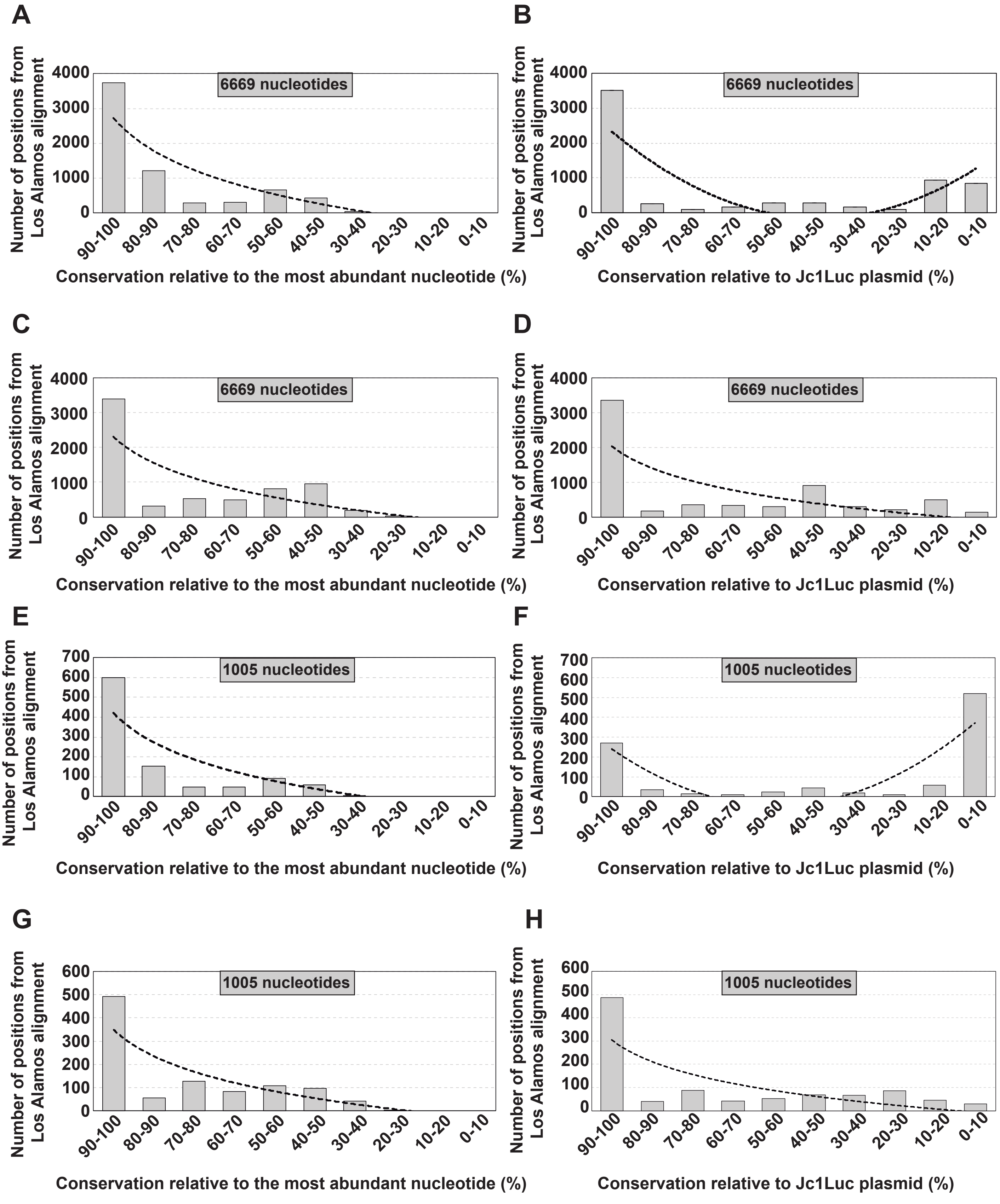
A
6669 nucleotides
Number of positions from Los Alamos alignment
0
1000
2000
3000
4000
90-100
80-90
70-80
60-70
50-60
40-50
30-40
20-30
10-20
0-10
Conservation relative to the most abundant nucleotide (%)
B
6669 nucleotides
Number of positions from Los Alamos alignment
Conservation relative to Jc1Luc plasmid (%)
C
6669 nucleotides
Number of positions from Los Alamos alignment
Conservation relative to the most abundant nucleotide (%)
D
6669 nucleotides
Number of positions from Los Alamos alignment
Conservation relative to Jc1Luc plasmid (%)
E
1005 nucleotides
Number of positions from Los Alamos alignment
0
100
200
300
400
500
600
700
Conservation relative to the most abundant nucleotide (%)
F
1005 nucleotides
Number of positions from Los Alamos alignment
Conservation relative to Jc1Luc plasmid (%)
G
1005 nucleotides
Number of positions from Los Alamos alignment
0
100
200
300
400
500
600
Conservation relative to the most abundant nucleotide (%)
H
1005 nucleotides
Number of positions from Los Alamos alignment
Conservation relative to Jc1Luc plasmid (%)

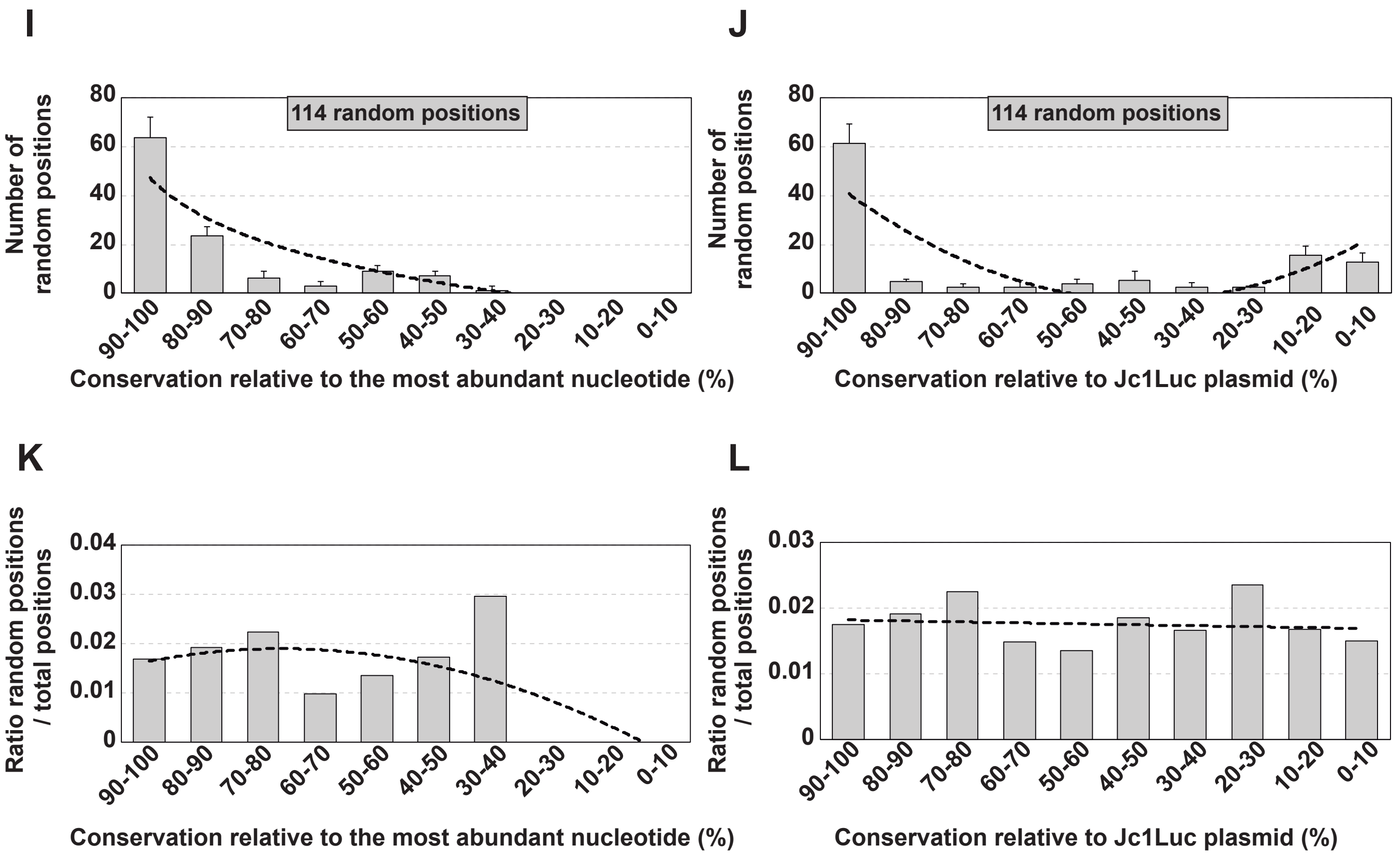

I
J
K
L
114 random positions
Number of random positions
Conservation relative to the most abundant nucleotide (%)
Conservation relative to Jc1Luc plasmid (%)
Ratio random positions / total positions
90-100
80-90
70-80
60-70
50-60
40-50
30-40
20-30
10-20
0-10

**Figure S3.** Resampling with specific genotypes, and simulations of distribution of HCV genomic nucleotides among conservation groups according to the LANL sequence alignment. **(A)** The positions comprised between the beginning of the NS2-coding region and the end of the NS5B-coding region (residues 2769 to 9377; H77 numbering) of an alignment of the HCV G1 and G2 genotypes (1112 sequences) were distributed in different windows according to the conservation of the most abundant nucleotide in each position of the Los Alamos alignment. Conservation groups are indicated in abscissa, and the number of positions that belongs to each group is indicated in ordinate. The discontinuous line corresponds to function $y = -1363\ln(x) + 2726.3$ ($R^2 = 0.7598$). **(B)** The positions were distributed in different windows according to the conservation of the nucleotide in the in the Los Alamos alignment, calculated using the HCV sequence in plasmid Jc1FLAG2(p7-nsGLuc2A) as reference. Conservation groups are indicated in abscissa, and the number of positions that belongs to each group is indicated in ordinate. The discontinuous line corresponds to function $y = 94.405x^2 - 1156x + 3390.2$ ($R^2 = 0.5997$). **(C, D)** Same as A and B except that conservation in the Los Alamos alignment was calculated using the same number of sequences of G1 and G2 (129 sequences of each genotype). The defining functions are C: $y = -1079\ln(x) + 2297.3$ ($R^2 = 0.6065$); D: $y = -910.9\ln(x) + 2042.8$ ($R^2 = 0.4731$). **(E, F)** Same as A, B except that the residues distributed among conservation groups are those of the NS5A-NS5B-coding region (residues 7584 to 8588; H77 numbering) for G1, G2, G3 and G4 genotypes (1191 sequences). The defining functions are E: $y = -212.8\ln(x) + 421.91$ ($R^2 = 0.7329$); F: $y = 17.186x^2 – 174.47x + 398.42$ ($R^2 = 0.695$). **(G, H)** Same as E, F except that 28 sequences of each of genotypes G1, G2, G3 and G4 were used for the alignment. The defining functions are G: $y = -164.6\ln(x) + 349.15$ ($R^2 = 0.6864$); H: $y = -135\ln(x) + 304.37$ ($R^2 = 0.5198$). **(I)** Triplicate simulation (using program Excel 2016) of the distribution of 114 random positions (within residues 2769 to 9377 (H77 numbering) (the same number of heterogeneity sites found in the HCV cell culture quasispecies) among conservation ranges calculated relative to the conservation of the most abundant nucleotide in the Los Alamos alignment. Conservation group are indicated in abscissa. The average and corresponding standard deviations of random positions is indicated in ordinate. The discontinuous line corresponds to function $y = -23.71\ln(x) + 47.219$ ($R^2 = 0.7806$). **(J)** Same as I except that conservation in the Los Alamos alignment was calculated relative to the HCV sequence in plasmid Jc1FLAG2(p7-nsGLuc2A). The discontinuous line corresponds to function $y = 1.6187x^2 – 19.999x + 59.078$ ($R^2 = 0.601$). **(K)** Ratio of the average of random positions that belong to each range of conservation group (calculated relative to the most abundant nucleotide as in panel I to the number of nucleotide positions of each group. The discontinuous line corresponds to function $y = -0.0005x^2 + 0.0031x + 0.0138$ ($R^2 = 0.4824$). **(L)** Ratio of the average of random positions that belong to each range of conservation group (calculated relative to corresponding nucleotide in the reference sequence Jc1FLAG2(p7-nsGLuc2A) as in panel J to the total nucleotide positions of each group. The discontinuous line corresponds to function $y = 0.0183e^{-0.008x}$ ($R^2 = 0.0187$). The random positions used for these controls are given in Table S4.
